# Interplay between material properties and cellular effects drives distinct pattern of interaction of graphene oxide with cancer and non-cancer cells

**DOI:** 10.1101/2024.11.01.621541

**Authors:** Yingxian Chen, Vinicio Rosano, Neus Lozano, YuYoung Shin, Aleksandr Mironov, David Spiller, Cinzia Casiraghi, Kostas Kostarelos, Sandra Vranic

## Abstract

Understanding how graphene oxide (GO) interacts with cells is crucial for its safe and efficient biomedical applications. Despite extensive research, a systematic investigation using a panel of cell lines, thoroughly characterized label-free nanomaterials, and complementary analytical techniques is lacking. Here, we examined the uptake of thin GO sheets with distinct lateral dimensions in 13 cell lines: 8 cancer (HeLa, A549, PC3, DU-145, LNCaP, SW-480, SH-SY5Y, U87-MG) and 5 non-cancer (BEAS-2B, NIH/3T3, PNT-2, HaCaT, 293T), using confocal microscopy, transmission electron microscopy, and flow cytometry. Our results reveal a striking difference in GO uptake: non-cancer cells internalized GO efficiently, while in cancer cells, GO predominantly interacted with the plasma membrane, showing minimal to no internalization. Comparison to other nanomaterials (polystyrene beads and graphene flakes) confirmed that cancer cells internalize materials similarly to non-cancer cells, indicating GO-specific interactions. We identified that GO thickness plays a pivotal role in this differential uptake. Additionally, GO disrupted the actin cytoskeleton, impairing migration in cancer but not in non-cancer cells. We propose that thin GO sheets act as a cue upon interaction with the plasma membrane of cancer cell lines, subsequently inducing actin filaments disruption leading to impaired endocytosis, migration activity, and reduced capacity of cancer cells towards GO uptake.

## Introduction

Graphene oxide (GO), due to aqueous dispersibility, large surface area suitable for the loading of therapeutic cargos through surface functionalisation and excellent biocompatibility has received extensive research interest for various biomedical applications^1–3^. However, to fully explore the biomedical potential of GO and assure its safe use, it is crucial to understand how GO interacts with biological systems at the cellular level.

Currently, a considerable number of *in vitro* studies have investigated interactions and uptake profile of GO. These studies mostly focused on cancer^4–26^, normal^27–32^, or immune cells^33–35^, with the vast majority of the studies using immortalised, commercially available cell lines. Considering the surging interest in using GO in cancer therapy, especially as drug carriers, it is critical to understand the uptake pattern of GO in cancer and normal cells. Some studies have compared the uptake of GO between cancer and normal cells^36–39^, cancer and immune cells^40–43^, or a combination of cancer, normal, and immune cells^44–46^. However, these studies largely used only one of each cell type^36,40,41,44^ or a total of no more than three cell lines for comparison^38,39,42,43,46^. One study examined the uptake of GO in six cell lines (three cancer, one normal, and two immune cell lines) and reported uptake of GO only in immune cells^45^. Thus, no existing studies have compared a large panel of cancer and normal cells, where commonalities and differences between GO interaction with diseased and healthy cells could be explored in-depth. Studies reporting GO uptake in cancer and normal cells have been contradicting^36–39,44–46^. Some studies showed the uptake of GO in both cancer and normal cells^38,39,44,46^, and others showed no uptake of GO in cancer and/or normal cells^36,37,45^. In some cases, GO is modified with different moieties to enhance cancer cell interaction compared to normal cells^37,39^. One factor that may have added to the inconsistent findings is that some studies involved the attachment of labels to enable GO tracing and uptake detection in cells^36,38,46^. Consequently, the intrinsic properties of GO can be altered by the labelling process, and the risk of misinterpretation due to potential label detachment is also increased. Furthermore, the selection of analytical techniques for the uptake assessment of GO is critical. Confocal laser scanning microscope (CLSM), transmission electron microscopy (TEM), and flow cytometry are three of the most common techniques used for cellular uptake assessments. However, each of these techniques has its advantages and shortcomings. Briefly, CLSM allows efficient distinguishment between internalised and cell surface adsorbed material in a time-efficient manner, but fluorescent labelling of material is often needed. TEM is an extremely powerful technique that offers high-resolution images, but it is a low throughput technique, so assessment by TEM is often restricted to a small sample size of cells. Also, considering that cellular compartments and GO are built mainly of carbon atoms, clear distinguishment between the material and cellular compartments is not always easily achieved. Finally, flow cytometry quantifies the total cellular interaction of material but cannot distinguish between internalised material and those adsorbed on cells. Thus, CLSM, TEM, and flow cytometry are good complementary techniques, but ambiguity could arise due to over-reliance on only one of these techniques, particularly flow cytometry. However, surprisingly, flow cytometry has been frequently used as a stand-alone or main analytical technique for assessing cellular uptake of GO^20,21,33,42–46^, leading to contradictory findings. For example, while both Yue et al. and Paino et al. investigated the uptake of non-functionalised label-free GO (with similar sizes) by flow cytometry, the former reported that cancer and normal cells (MCF-7, Hep G2, LLC, and HUVEC) were not able to internalise the material, but the latter reported the uptake of GO in both cancer and normal cells (HeLa, L929)^44,45^.

To close the identified research gap, we systematically compared the uptake of label-free, non-functionalised GO in a panel of cancer and normal cell lines combining CLSM, flow cytometry and TEM. Furthermore, we identified the properties of GO and cellular effects that influence the way GO interacts with cancer and non-cancer cells.

## Result and discussion

### Interactions of GO with cancer and non-cancer cells

We first assessed the cellular uptake of GO by confocal imaging. We used thoroughly characterized small-GO (s-GO, 25 nm – 2.4 µm) and ultrasmall-GO (us-GO, 10 – 790 nm) in the present study (**Figure S1**, **Table S1**). **Figure S2** shows the interaction of GO (s- and us-GO, 50 μg/mL) in NIH/3T3 (non-cancer) and HeLa (cancer) cells after 24 h of incubation by CLSM. Cellular uptake of GO is indicated by the localisation of the GO signal (in red) within the plasma membrane (stained green); internalised GO is shown as concentrated red spots localised towards the centre of the cell. On the other hand, extracellular GO interacting with the plasma membrane is shown as dispersed clouds of red signals surrounding or localised on top of cells. Interestingly, we identified two distinct GO interaction patterns between the two cell lines. In NIH/3T3, both s- and us-GO were found internalised and interacting with the plasma membrane, with no obvious differences in the uptake quantity. Meanwhile, for HeLa cells, both s- and us-GO were found to interact with the plasma membrane, and no intracellular distribution of the materials was observed. This pattern agrees with the results from our previous study, where we observed the uptake of GO in the non-cancer cell lines^47^. The uptake of GO (s- and us-GO at 25, 50, and 75 μg/mL for 24 h) was then assessed in a total of eight cancer and five non-cancer cell lines by CLSM (**Figure 1**, **S3–S14**, and **Table S2**). Surprisingly, the non-cancer cells were repeatedly found to take up both s-GO and us-GO, but none of the cancer cell lines were found to take up the two materials. Further analysis in cancer (HeLa, LNCap, PC3) and non-cancer (BEAS-2B, NIH/3T3, PNT-2) cells showed the same interaction pattern remained for large-GO (l-GO, 1.5 – 25.5 µm) (**Figure S15**).

**Figure 1:**
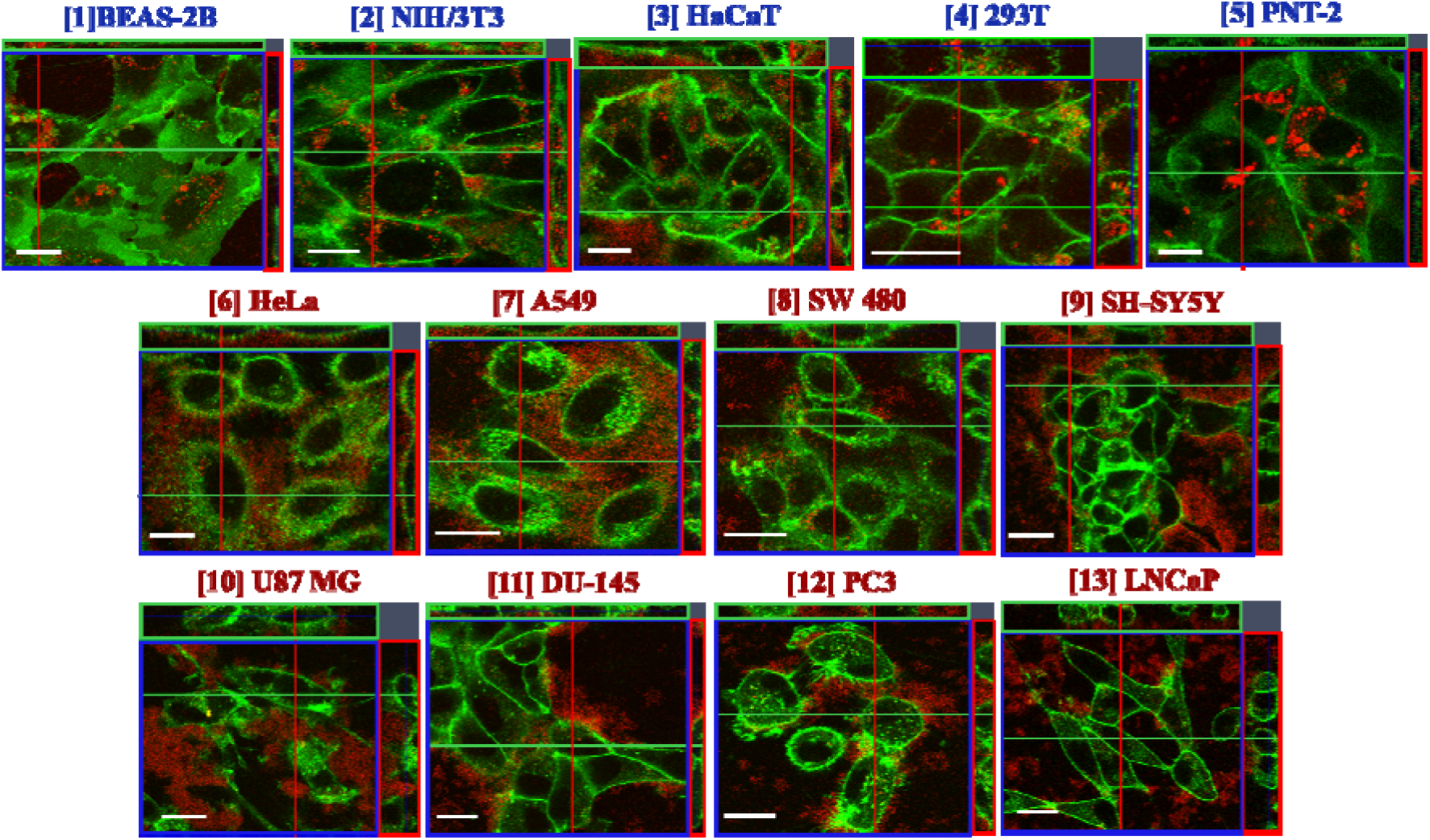
Interaction of GO (s- and us-GO) in cancer and non-cancer cells, assessed by CLSM (orthogonal projections of the middle section of the cell are shown). Non-cancer cells: **[1]** BEAS-2B (s-GO treated), **[2]** NIH/3T3 (us-GO treated), **[3]** HaCaT (s-GO treated), **[4]** 293T (us-GO treated), **[5]** PNT-2 (s-GO treated). Cancer cells: **[6]** HeLa (s-GO treated), **[7]** A549 (us-GO treated), **[8]** SW 480 (s-GO treated), **[9]** SH-SY5Y (us-GO treated), **[10]** U87 MG (s-GO treated), **[11]** DU-145 (us-GO treated), **[12]** PC3 (s-GO treated), and **[13]** LNCaP (s-GO treated). See **Figure S3 – 14** for the full panel of GO (s- and us-GO) interaction profiles in cell lines **[2]** – **[13]**, respectively. See Chen et al.^47^ for in-depth GO (s- and us-GO) interaction profiles and uptake mechanisms in **[1]** BEAS-2B cells. See **Table S2** for the summary of GO (s- and us-GO) interaction profiles in all cell lines. Green = plasma membrane, Red = GO. Scale bar = 20 µm.

All studied cell lines were maintained in cell-type specific media either recommended by the American Type Culture Collection (ATCC) or previously determined experimentally. However, in all GO treatments (unless specified), the same cell culture media (RPMI-1640 with 10% FBS) was used to exclude variability induced by different media compositions. To examine the effect of treatment media on the uptake of GO, we compared the uptake of GO in cancer cells under different treatment media. In human epithelial prostate carcinoma cells (DU 145), we observed no apparent changes in the uptake profile of GO (s- or us-GO) using RPMI-1640 (see **Figure S12a**) or DU 145 cells specific media (see **Figure S12b**). In agreement, human epithelial glioblastoma cells (U87 MG) also showed comparable GO interaction patterns using RPMI-1640 (see **Figure S11**) or U87 MG cell-specific media^48^. Therefore, the choice of treatment media was not the dominant factor driving the uptake difference of GO between cancer and non-cancer cells.

#### Assessment by TEM

To confirm the observations by CLSM, we next examined the uptake of GO using TEM (**Figure 2a – d**). As expected, internalised GO was found within the vesicles of BEAS-2B and NIH/3T3 cells, with no obvious signs of cytosol localised GO flakes (**Figure 2a – b**). In contrast, most HeLa and A549 cells showed no sign of GO internalisation (**Figure 2c – d**). Thus, this finding confirmed the uptake difference of GO in cancer and non-cancer cells, as identified by CLSM. However, we noticed that cancer cells attempted to take up the materials, as indicated by plasma membrane ruffling and deformations near GO (**Figure S16**). Also, although most of the s- and us-GO were found interacting with the cancer cell plasma membrane without being taken up, very few cancer cells were found with internalised small amounts of us-GO, localised within intracellular vesicles (**Figure S16**).

**Figure 2:**
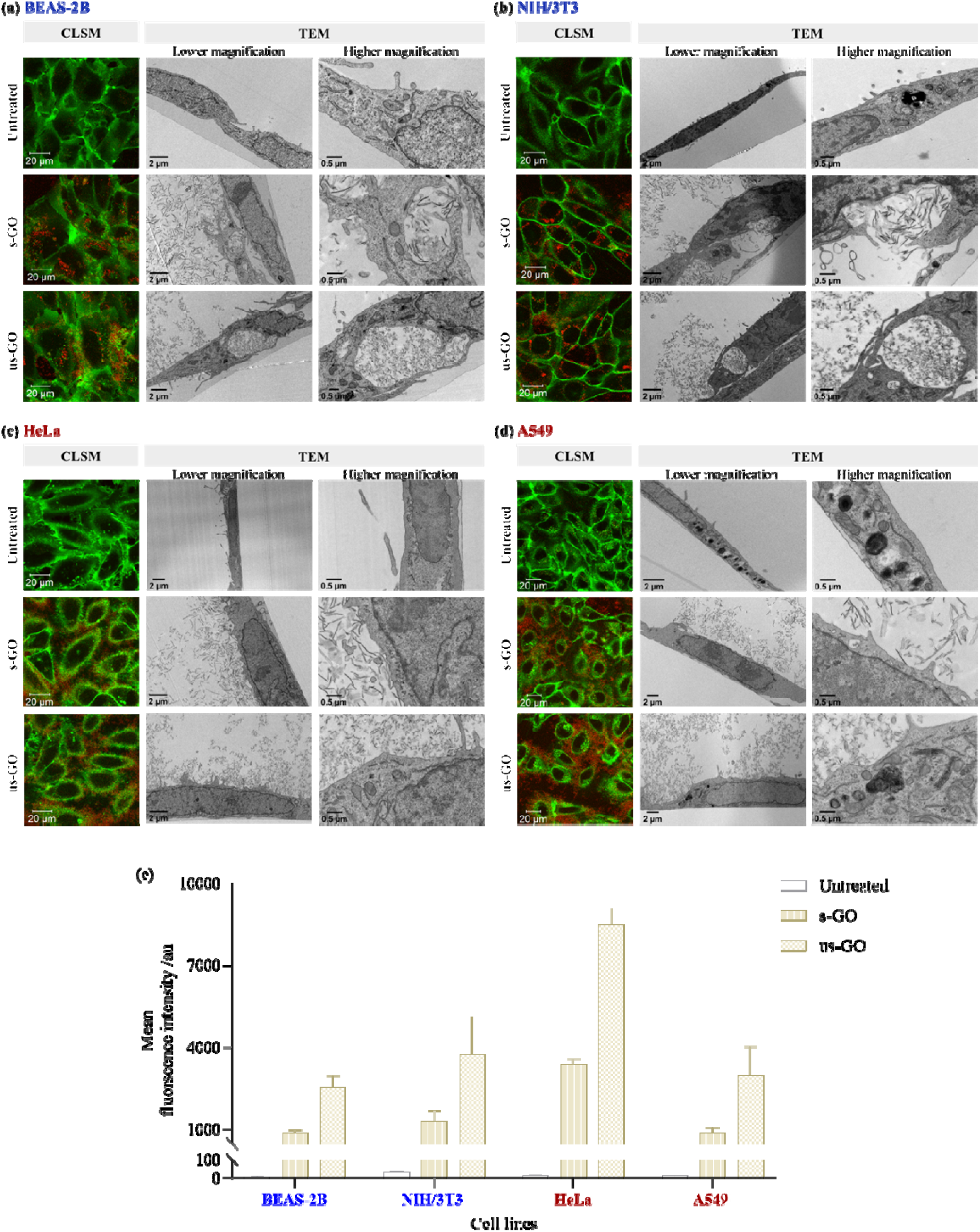
Interaction of GO (s- and us-GO, 50 μg/mL) with **(a)** BEAS-2B, **(b)** NIH/3T3, **(c)** HeLa and **(d)** A549 cells by CLSM, TEM, and **(e)** flow cytometry. For analysis by flow cytometry, the cells were collected with PBS (−/−) washing; see **Figure S17** for cells collected without PBS (−/−) washing. See **Table S3** for the corresponding statistical analysis of the interaction of GO with cells by flow cytometry. Green = plasma membrane, red = GO.

#### Assessment by flow cytometry

We further examined the cellular interaction of GO using flow cytometry, which quantifies the total interaction of GO with cells (i.e., the sum of GO taken up and adhered to the cells). The interaction of GO (s- and us-GO) with cancer (HeLa, A549) and non-cancer (BEAS-2B, NIH/3T3) cell lines were compared, with (**Figure 2e**) and without (**Figure S17**) washing prior to flow cytometry examination. For both cancer and non-cancer cell lines, washing greatly reduced the mean fluorescent intensity (MFI) of GO (s- and us-GO), which confirmed the presence of GO bound to the cell surface (see **Figures 2e** and **S17**). However, regardless of the types of GO and whether washing was performed, the MFI for HeLa cells was significantly higher than for the other three cell lines. In contrast, no significant differences were observed between A549, BEAS-2B and NIH/3T3 cells (see **Table S3a** and **Figure S17** [**table a**]).

As both HeLa and A549 cells clearly showed minimal uptake of GO by CLSM and TEM, we concluded that GO adhered to the surface of the cells contributed to the fluorescent signal measured by flow cytometry. The difference in MFI between HeLa and A549 cells demonstrates that GO adhered more strongly to the HeLa cell membrane than the A549 cell membrane (**Figure 2e**). Furthermore, the MFI for us-GO was significantly higher than s-GO in all studied cell lines, with and without washing (see **Table S3b** and **Figure S17** [**table b**]). Considering the spectrofluorometric analysis of the emission spectra of GO showed that the intrinsic fluorescent intensity of s-GO was higher than us-GO (**Figure S18**), the finding suggests that us-GO was better adsorbed to the plasma membrane of the cells than s-GO. Previous studies have reported the interaction of non-functionalised label-free GO with cancer cells using flow cytometry^20,21,44^. Several studies concluded on the uptake of GO in A549 cells^20,21^, using flow cytometry^42–44,46^. Thus, it is important to recognise that uncertainty could arise from the presence of extracellular bound materials in uptake studies mainly based on flow cytometry.

Likewise, the uptake of GO in cancer cells by TEM has been widely reported^5,16,22,23,40^. However, it is essential to acknowledge that the cell sample size analysed by TEM is relatively small compared to other techniques such as CLSM and flow cytometry. Therefore, uptake studies solely based on TEM can suffer from the overemphasis on the success or failure of the uptake but are rarely concerned with the level and scale of uptake.

Proceeding, our results showing successful uptake of GO in non-cancer cells are in agreement with the previous findings^28,32,36,47^. However, the uptake of GO in non-cancer cells has been shown to depend on the cell phenotype^31^. For example, Kucki et al. showed that GO was internalised by undifferentiated intestinal cells but not the differentiated intestinal cells with hair-like surface structures^31^. Thus, cell surface topography that permits better association of GO with the plasma membrane allows enhanced GO uptake.

### Assessment of the uptake capacity of cancer and non-cancer cells

Size, shape, and surface chemistry are the most extensively studied physico-chemical properties critical for NMs-cellular interaction. To explain the observed differential uptake of GO by cancer and non-cancer cells, we interrogated the capacity of cancer and non-cancer cells towards the uptake of NMs with different size, shape, and surface chemistry. The NMs used were divided into two groups: **(1)** commercially available polystyrene microspheres with different size and surface charge (negatively charged carboxylate-modified beads [0.1, 0.5, and 1 μm] and positively charged amine-modified beads [0.2 and 1 μm]), and **(2)** graphene flakes with different surface charge (negatively charged 1-pyrenesulfonic acid sodium salt stabilised graphene dispersion [Gr-PS1] and positively charged trimethylammonium linked pyrene stabilised graphene dispersion [Gr-TMA_3_]). produced and characterized as already described^30,49,50^.

Microspheres were selected as spheres are one of the most commonly used types of NMs^51^, and they are typically better internalised than two-dimensional (2D) NMs^36^. By interrogating cellular capacity to take up microspheres of different sizes and functionality, we aimed to confirm if cancer cells, in general, have a lower endocytic ability than non-cancer cells or if this applies only to materials with specific properties. Second, using graphene flakes of different functionalities and surface charges, we aimed to confirm the ability of cancer cells to internalise other 2D NMs.

#### Uptake of microspheres

**Figures 3** and **4** show the interaction of cancer and non-cancer cells with the FITC-labelled microspheres with positive (+) and negative (−) surface charge, assessed by CLSM and flow cytometry, respectively. **Figures 3** and **4** showed that for both cancer and non-cancer cells, the uptake of the −1 μm beads was comparable to the +1 μm beads. However, BEAS-2B cells exhibited the highest uptake, followed by HeLa, NIH/3T3 and A549 cells as evidenced by MFI values: BEAS-2B (−1 μm = 20000, +1 μm = 18871), NIH/3T3 (−1 μm = 4735, +1 μm = 5174), HeLa (−1 μm = 5756, +1 μm = 6438) and A549 (−1 μm = 4270, +1 μm = 4531). Likewise, the uptake of −0.5 and −0.1 μm beads in cancer and non-cancer cells exhibited a similar trend to the 1 μm beads: the MFI of the beads was highest in BEAS-2B cells (−0.5 μm = 6341, −0.1 μm = 1843), followed by NIH/3T3 (−0.5 μm = 2168, −0.1 μm = 662), HeLa (−0.5 μm = 1545, −0.1 μm = 545) and A549 (−0.5 μm = 866, −0.1 μm = 344). On the other hand, the highest uptake of the +0.2 μm beads was observed in HeLa cells (22614), followed by BEAS-2B (12707), A549 (11472) and NIH/3T3 (8356) cells (see **Figures 3** and **4**).

**Figure 3:**
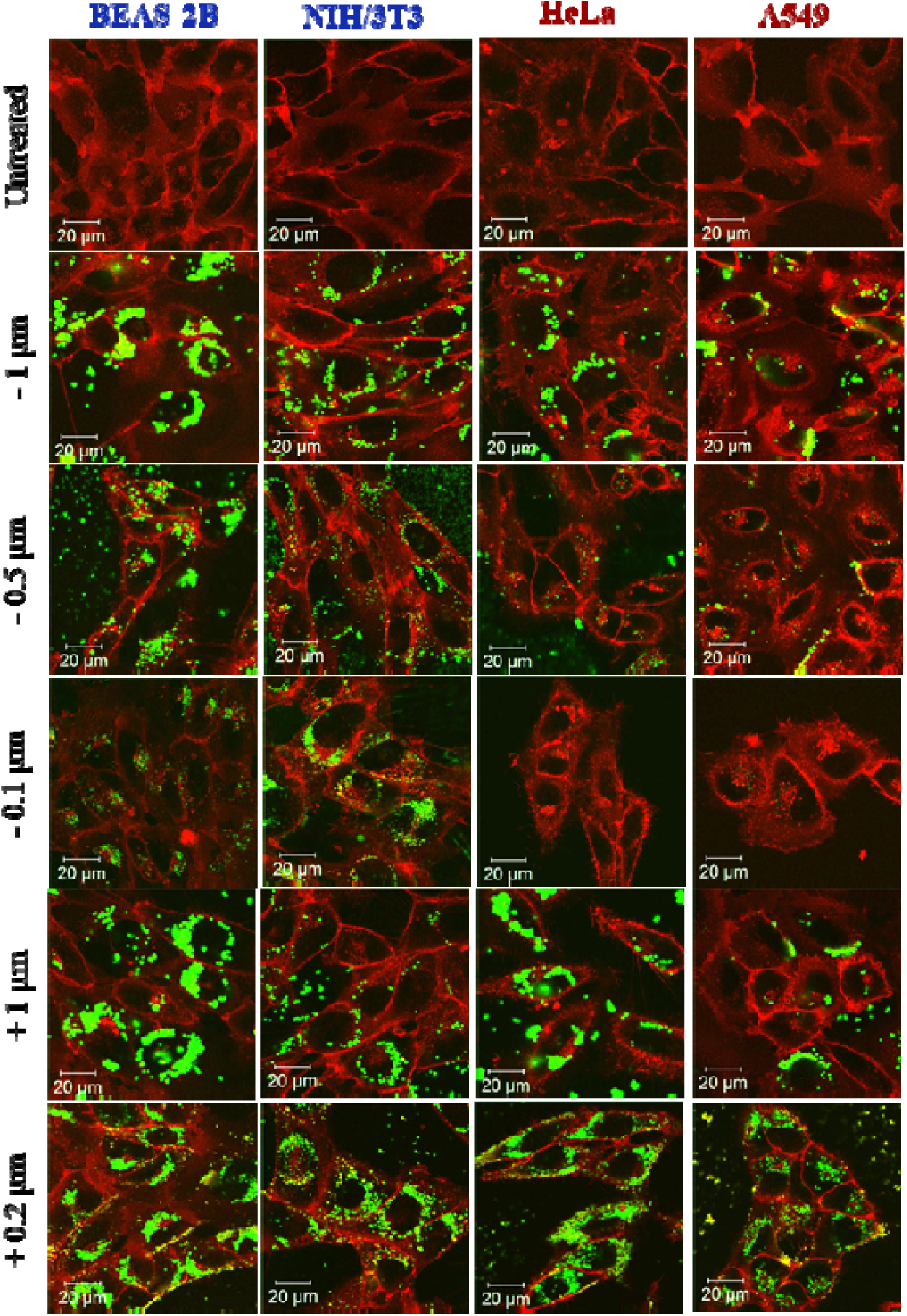
Interaction of BEAS-2B, NIH/3T3, HeLa and A549 cells with negatively charged (−) 0.1, 0.5, and 1 μm carboxylate-modified, and positively charged (+) 0.2 and 1 μm amine-modified beads by CLSM. Red = plasma membrane, green = FITC-labelled beads. Scale bar = 20 μm.

**Figure 4:**
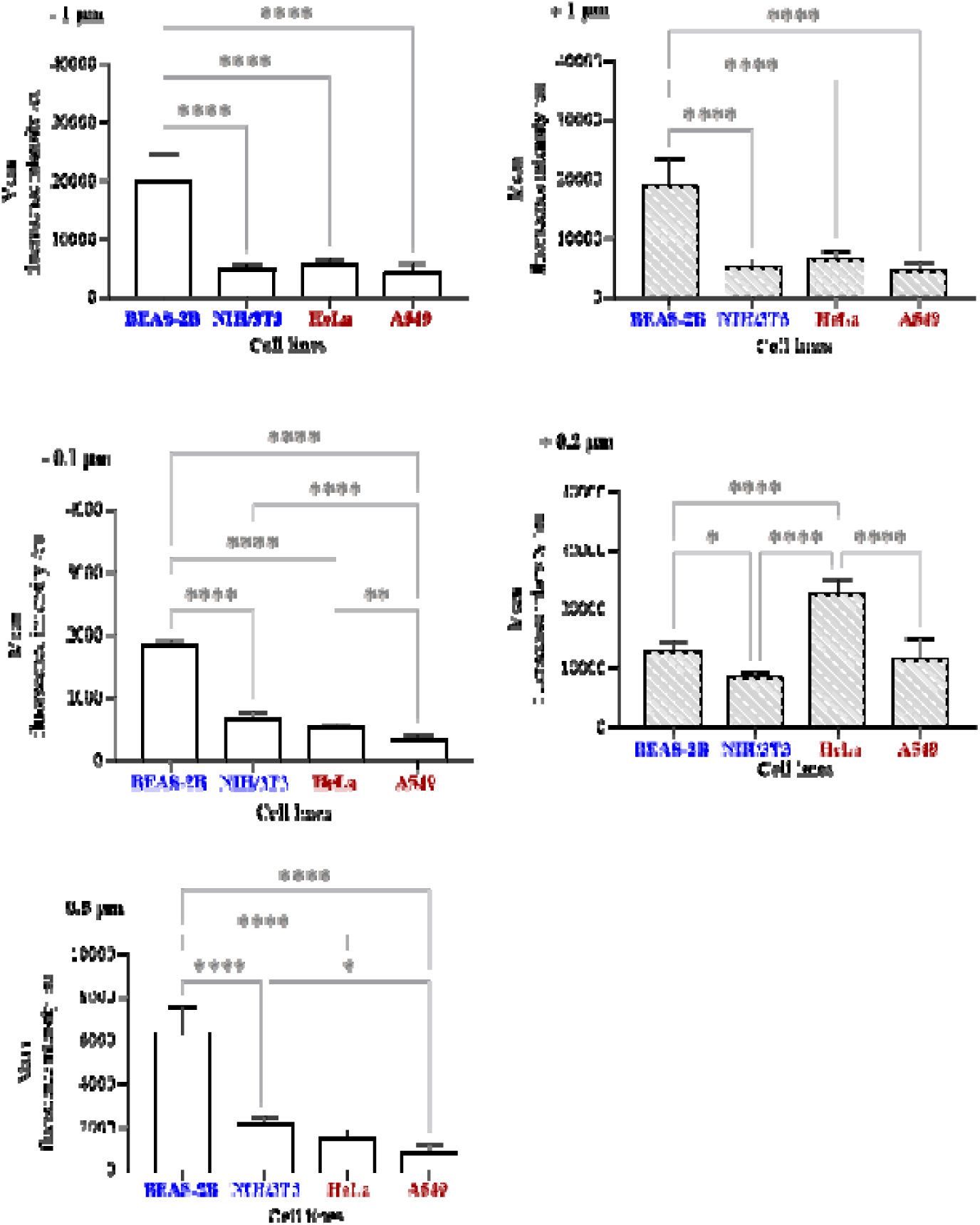
Interaction of BEAS-2B, NIH/3T3, HeLa and A549 with negatively charged (−) 0.1, 0.5, and 1 μm carboxylate-modified, and positively charged (+) 0.2 and 1 μm amine-modified beads by flow cytometry. The data were statistically analysed using analysis of variance (one-way ANOVA) with Tukey’s multiple comparisons test. n = 3 with duplicates. *Statistically different: *\*p* < 0.05, *\*\*p* < 0.01 and *\*\*\*\*p* < 0.0001.

Despite variability in the level of uptake, both cancer and non-cancer cells could take up all the beads irrespective of their size or charge. Furthermore, the fact that the NIH/3T3 cell showed no significant difference in the uptake of −1 μm beads compared to the A549 and HeLa cells demonstrates that cancer cells do not, intrinsically, exhibit a lower uptake capacity towards negatively charged materials. This result is further supported by the comparable uptake of −1 and +1 μm beads in each cell line. Both cancer and non-cancer cells were found to take up the small positively charged (+0.2 μm) beads better than the small negatively charged (−0.1 μm) beads. Yet, it is important to note that of all bead types, the +0.2 μm beads were also most uptaken in HeLa, A549, and NIH/3T3 cells. Hence, the result suggests that the role of surface charge on the uptake of material may become more significant for the material below a certain size.

#### Uptake of Graphene flakes

We next investigated the uptake of Gr-TMA_3_ and Gr-PS1 in cancer (HeLa) and non-cancer (BEAS-2B) cells. As shown in **Figure S19**, due to the light absorption properties of graphene flakes, the material appears black in the bright field channel. On the other hand, the cell was stained green with the fluorescein diacetate (FDA) dye, and where the cell had taken up the material, the flakes quenched the signal of the dye (see **Figure S19**). It is apparent from **Figure S19** that both Gr-TMA_3_ and Gr-PS1 were internalised in BEAS-2B and HeLa cells, which confirms that cancer cells could take up positively or negatively charged 2D NMs. However, in both BEAS-2B and HeLa cells, the positively charged Gr-TMA_3_ were internalised better than the negatively charged Gr-PS1 flakes. Considering that graphene flakes had an average size of approximately 200 nm^30^, this is in line with results from the uptake study of the microspheres, where the small positively charged microspheres were taken up better than the small negatively charged ones (see **Figures 3** and **4**). In addition, BEAS-2B cells were noticeably taking up Gr-PS1 better than HeLa cells (see **Figure S19**). This is also in line with those observed from the uptake study of the microspheres, where the small negatively charged microspheres were taken up better in BEAS-2B than HeLa cells (see **Figures 3** and **4**).

Findings from the uptake study of microspheres and graphene flakes demonstrate that the material surface charge affects its uptake in cancer and non-cancer cells, but not in an “all or nothing” manner. As is well-known, the material surface charge affects the electrostatic interactions between the material and plasma membrane^52^. Positively charged NMs have been reported to internalize better than their negatively charged counterparts due to a more energetically favourable interaction with the cell surface that enhances the endocytosis process^52,53^. Indeed, it has been suggested that the lack of GO uptake in non-phagocytic cells is due to the electrostatic repulsion between the material and plasma membrane^45^. However, we observed GO interaction with both cancer and non-cancer cell plasma membranes, and our result by flow cytometry demonstrate that GO adhered strongly to the cancer cell surface (**Figures 1**, **2**, and **S2–17**). In fact, the adhesion of GO on the plasma membrane of cancer cells has been widely acknowledged by different research groups^21,24,32,41^.

It is worth noting that GO may interact more with the cancer cell plasma membrane than the microspheres/graphene flakes. As shown by CLSM (**Figure S2**, the middle section of cells), GO was displayed as a uniform cloud of signal surrounding the HeLa cell plasma membrane, outlining the shape of HeLa cells. In comparison, substantially fewer microspheres/graphene flakes were found surrounding the HeLa cell plasma membrane (**Figures 3** and **S19**). The difference in material thickness may have contributed to the difference between GO and the microspheres/graphene flake in the plasma membrane interaction. Being mostly monolayer, over 90 % of the GO sheet had a thickness of 2 nm or less (see **Table S1**). In contrast, while the microspheres had a thickness range of 0.2 - 1 μm, the graphene flakes existed mostly as few-layers graphene (70 and 61 % for Gr-TMA_3_ and Gr-PS1, respectively) with a thickness range of 2 – 20 nm^30,49,50^.

Previous research into the effect of Gr thickness by simulation showed that few-layer graphene (3 or 9 layers) exhibited an increase in the energy barrier for piecing the lipid bilayer compared to the single-layer graphene^54^. It can be expected that thickness influences the specific surface area by volume of the material, and these differences can directly affect the colloidal behaviour of the material, which ultimately results in the formation of different biological interfaces when interacting with cells^55^. Thus, despite its two-dimensionality, the colloidal behaviour of graphene flakes may share commonality with the three-dimensional microspheres, contributing to a more similar pattern of cellular interaction. In contrast, the thinner GO possess greater two-dimensionality and remains as individualised 2D sheets when interacting with cells, as shown by TEM (see **Figure 2**).

Taken together, we clearly demonstrated that cancer cells take up materials with diverse properties, including shape, size, or surface charges comparable to GO. However, neither the uptake of microspheres nor the graphene flake occurs in the cell-type specific manner as observed for GO. We therefore conclude that the difference in thickness between GO and microsphere/graphene flakes contributes to the cell-type specific uptake of GO.

### Assessment of the effects of GO on endocytosis-relevant biological properties of the cells

#### The effect of GO on actin filaments

It was reported that GO could induce biological changes in cells, such as alteration of the plasma membrane composition^56^, cytoskeleton^13,20,41,57^ and migratory ability of cells^19,58^. Therefore, we asked if GO induces differential changes in endocytosis-relevant biological properties of cancer and non-cancer cells, ultimately contributing to the observed differences in GO uptake.

The effect of GO on cytoskeleton was examined by actin filament staining. Remodelling of the cytoskeleton is a prerequisite for endocytosis^59,60^. In particular, actin filaments are crucial for all three major uptake pathways^61^; actin filaments disruption agents such as latrunculin and cytochalasin D are well-known pharmaceutical inhibitors of endocytosis^62^.

**Figure 5** and **Figure S20** show the actin filament network for BEAS-2B and HeLa cells with and without GO (s- or us-GO, 100 μg/mL, 24 h), stained using the Alexa-Fluor 488-labelled phalloidin (**Figure S21** for NIH/3T3 and A549 cells). In addition, methyl-beta-cyclodextrin (M /J CD), which disrupts the cytoskeleton by depletion of the cholesterol in the cell membrane, was used as the positive control. Images in **Figures 5**, **S20** and **S21** are shown in the range indicator mode to better compare the actin filament network. The images are displayed in black and white, with red and blue areas. Red and blue represent areas with saturated and zero intensity, respectively. At the same time, black and white colours represent lower and higher intensity areas, respectively. The settings were adjusted to display the background in blue, and the same intensity settings were applied within each cell line.

**Figure 5:**
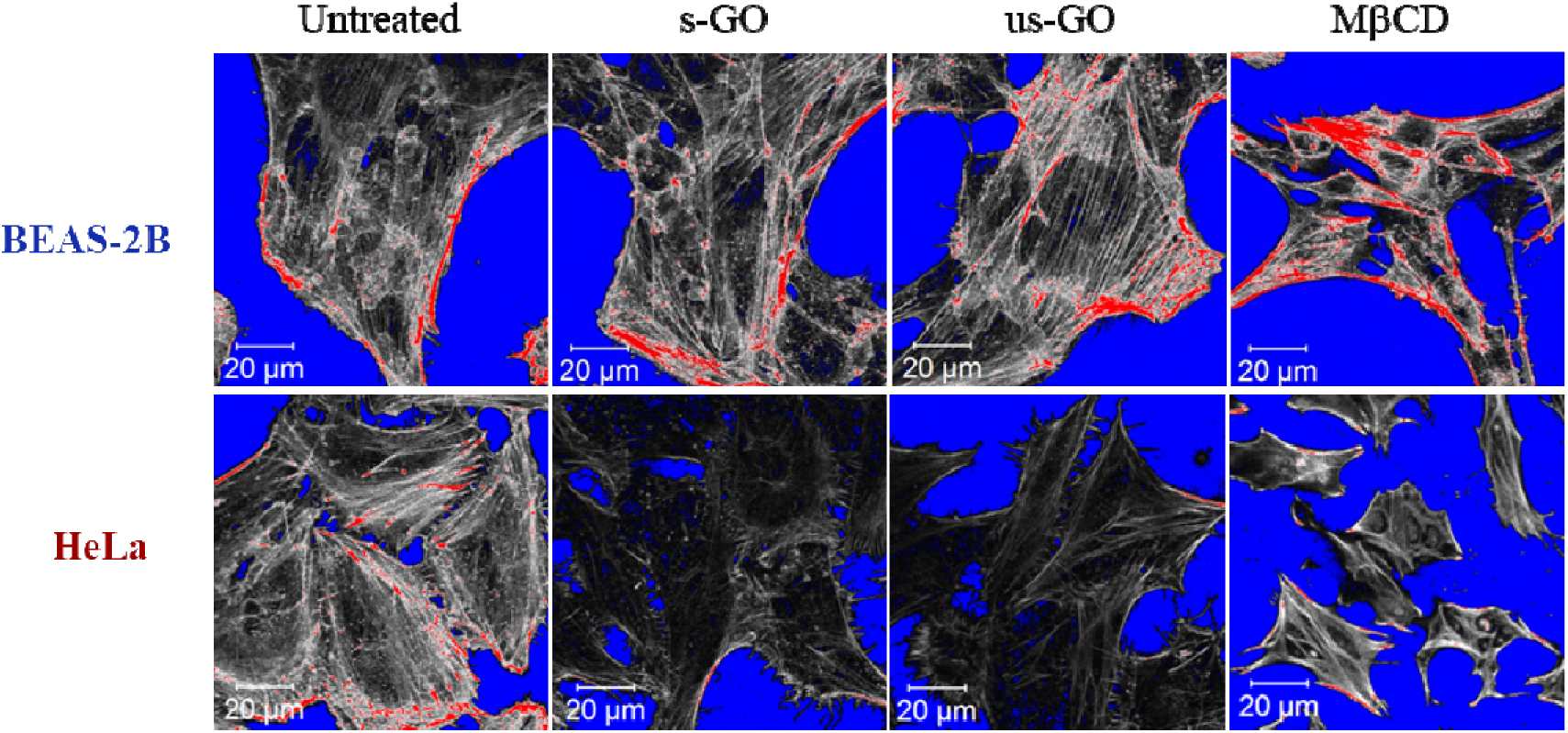
Staining of actin filament in BEAS-2B and HeLa cells, treated with and without GO (s-GO or us-GO at 100 μg/mL, 24 h) or MβCD (5 mM, 4 h, positive control). Cells were stained with the phalloidin 488 dye for actin filament, and the images are shown in the range indicator mode: red and blue represent saturated and zero intensity areas, whereas black and white represent lower and higher intensity areas, respectively.

For all four cell lines, it is apparent that M /JCD induced significant disruption to the actin filament network, resulting in an altered cell morphology compared to the untreated cells (see **Figures 5**, **S20** and **21**). In general, M /JCD treated cells appeared smaller than the untreated cells. In addition, features including the loss of the parallel distribution of the filaments, formation of actin spots, fragmented fibres, and increased stress fibres can also be identified in M /JCD treated cells. On the other hand, the result showed that GO (s- and us-GO) could induce changes to the actin filament network in cancer cells, but no apparent differences were noted in the non-cancer cells (see **Figures 5**, **S20** and **21**). One observed similarity in the effect of GO (s- and us-GO) on HeLa and A549 cells was that the actin filament density appears to be reduced in the GO treated cells compared to the untreated cells, shown by the increased lower intensity areas in GO treated cells (see **Figures 5**, **S20** and **21**). However, neither s-GO nor us-GO in HeLa or A549 cells could induce changes to a level comparable to M /JCD, where a pronounced reduction in the shape and size of cells was observed.

The actin cytoskeleton is heavily involved in many endocytic pathways, including clathrin-mediated endocytosis (CME), caveolae-mediated endocytosis (CavME) and macropinocytosis^62^. Notably, we previously confirmed GO internalisation in BEAS-2B cells *via* these three pathways^47^. Furthermore, disruption to the organisation and dynamic of the actin cytoskeleton has been widely recognised as the effect of various endocytosis inhibitors^62^. Hence, this result provides some tentative initial understanding of the observed GO uptake difference between cancer and non-cancer cells. Yet, the finding raised the intriguing question regarding the underlying mechanism for the remodelling of the actin cytoskeleton by GO and the selectivity toward cancer cells.

In agreement with the present results, previous studies have reported the reduction of the actin filament density in cancer cells with GO treatment^20,41^. It has been suggested that GO impairs the cytoskeleton by direct interaction^20,63^ or indirectly through suppression of the cell adhesion system^41^. Tian et al. and Bera et al. suggested the direct interference of GO with the actin filament and microtubule network in A549 and HCT116 cancer cells, respectively^20,63^. However, in these studies, the main evidence for GO’s direct interference with the cytoskeleton was based on GO incubation with the isolated actin filaments and tubulin protein. In our study, for cancer cells, GO was found predominantly surrounding the plasma membrane and not being available to directly interact with actin filaments in the cytosol.

In the study by Zhu et al., GO impaired the plasma membrane and lowered integrin expression in A549 cells, which subsequently affected the cell adhesion complex and the cytoskeleton network^41^. A growing body of evidence demonstrates that changes in cell biomechanical properties induce actin cytoskeleton reorganisation and modulate the uptake of nanoparticles^64–67^. For example, higher uptake of nanoparticles was observed for cells seeded on stiffer substrates^65,66^. Furthermore, substrate stiffness has been reported to regulate cell stiffness, with cell softening and reduced actin polymerisation observed for cells seeded on softer substrates^68^. Considering the growing amounts of evidence that shows cancer cells are softer than normal cells in 2D cultures^69^, and findings that show GO can modulate cancer cell mechanical properties by reducing cell stiffness and inducing actin cytoskeleton remodelling^70^, one possible hypothesis is that GO acts as mechanical cues that prompt further softening of cancer cells, resulting in cytoskeletal reorganisation, and ultimately affecting the endocytosis of GO.

#### The effect of GO on cell migration

Subsequently, the effect of GO on cell migration was investigated to validate its effect on actin filaments. Like endocytosis, regulation of the cytoskeleton network is essential for cell migration^71,72^. Therefore, the disruption of the actin cytoskeleton will affect cell migration. For instance, pharmaceutical inhibitors such as cytochalasin B, which disrupt actin filament polymerisation, inhibit cell migration as well as endocytosis^73,74^. We employed time-lapse tracking to assess the effect of GO on cell migration.

Herein, HeLa (one of the most studied cancer cell lines for GO cellular interaction) and BEAS-2B cells were used as representatives of cancer and non-cancer cell lines. BEAS-2B was chosen because we have previously established the uptake mechanism of GO in BEAS-2B cells^47^. In **Figure 6**, we compared migratory differences between untreated cells and cells treated with s-GO (50 μg/mL). Using cell trajectories, we observed that for both BEAS-2B and HeLa cells, untreated cells moved in a larger spreading area than cells treated with s-GO (see **Figure 6a**). However, the difference between the observed effect was greater for HeLa than BEAS-2B cells. This observation was further supported by the mean square displacement (MSD) measurement (see **Figures 6b** and **6e**), where the average surface area used by cells was plotted over time^75^.

**Figure 6:**
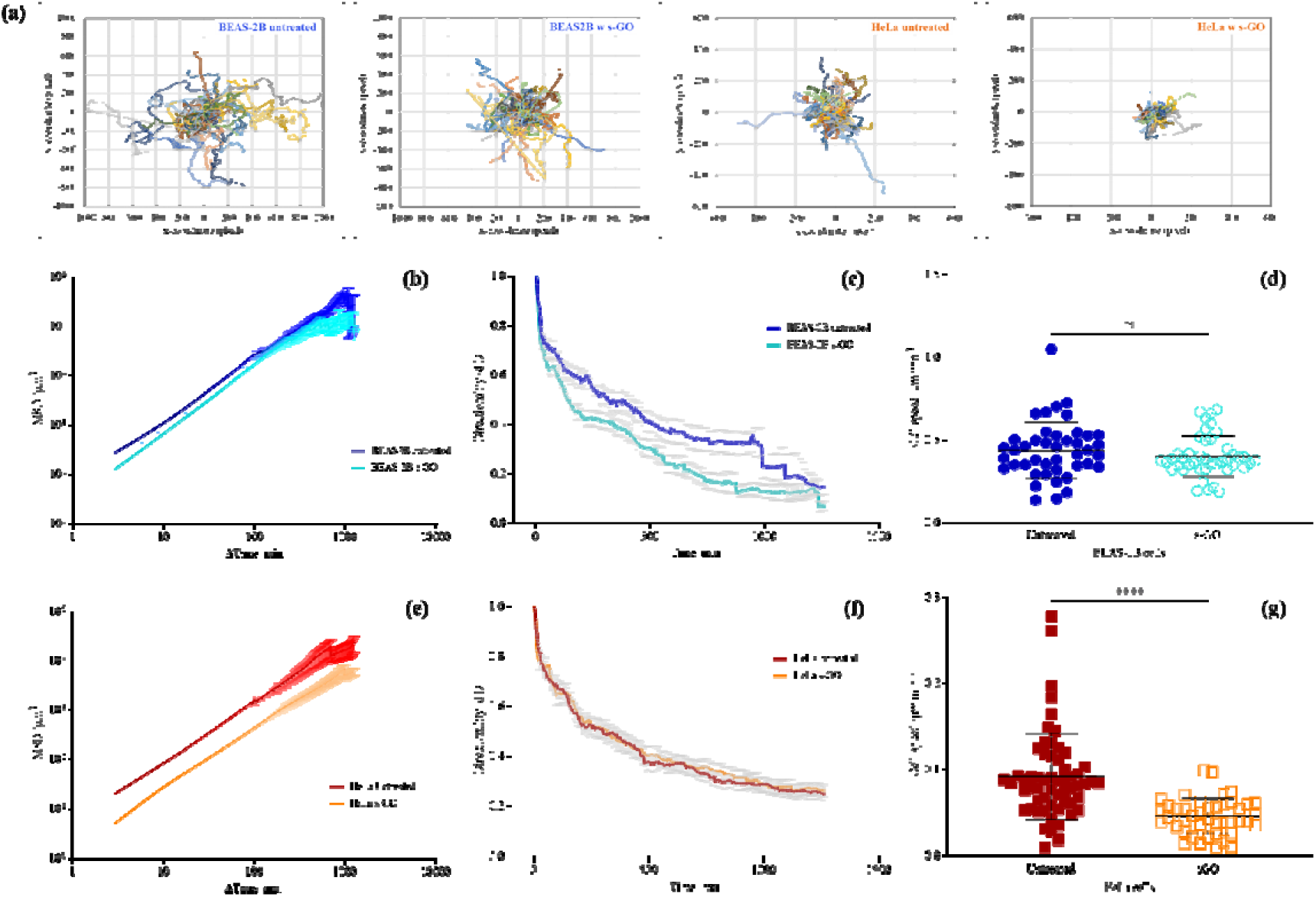
The effect of GO (50 μg/mL) on **(a** – **d)** BEAS-2B and **(a**, **e** – **g)** HeLa cell migration, assessed by live cell time-lapse tracking. Cell trajectories are visualised *via* **(a)** plot at the origins and further analysed *via* **(b**, **e)** MSD, **(c**, **f)** directionality analysis, and **(d**, **g)** cell speed measurement. Cell speeds are statistically analysed using an unpaired T-test. Number of cells analysed per condition = 43. *Statistically different: *\*\*\*\*p* < 0.0001, ns = not significant.

To understand better the effect of s-GO on BEAS-2B and HeLa cells, we further examined two key parameters that account for cell migration efficiency: directionality persistence and cell speed^75^. In **Figures 6c** and **6f**, the average directionality ratio over time is shown. The directionality ratio (d/D) for a set time point is the ratio of the straight-line distance (d) from the cell’s initial to end positions at the set time point over the actual distance (D) the cell travelled. Hence, if the directionality ratio equates to one, it suggests the cells have followed a perfectly straight path, whereas if the ratio approaches zero, it means the cells have followed a highly curved path. However, it should be noted that, in the absence of migration cues, directionality persistence for randomly moving cells will decay and vary considerably over time^75^. Therefore, instead of comparing the directionality ratio at one set time point (e.g., at the endpoint), assessing the directionality ratio over the time course better represents directional persistence.

As shown in **Figure 6c**, the BEAS-2B untreated cells followed a straighter path than s-GO treated BEAS-2B cells. This is shown by the rapid decay of the directionality curve in s-GO treated cells (mean d/D: 0.236 ± 0.008) compared to untreated cells (mean d/D: 0.402 ± 0.008). Whereas for HeLa untreated and s-GO treated cells, no apparent differences in the directionality ratio were observed over time (see **Figure 6f**). The directionality curve for HeLa untreated (mean d/D: 0.359 ± 0.006) and s-GO treated (mean d/D: 0.372 ± 0.006) cells stay overlapped over the entire time course of cell trajectory. On the other hand, the HeLa cell speed reduced significantly with s-GO treatment, but comparable cell speeds were found for BEAS-2B cells with or without s-GO treatment (see **Figure 6d** and **6g**, respectively). For example, the average speed for HeLa untreated cells was 0.091 ± 0.050 μm/min, which was reduced by 50.5 % in s-GO treated cells (0.045 ± 0.021 μm/min). In contrast, only an 8 % difference was found in the average speed between BEAS-2B untreated (0.438 ± 0.170 μm/min) and treated (0.403 ± 0.124 μm/min) cells.

The observed effect on the speed and spreading area of s-GO treated HeLa cells supports the finding that GO induced actin filament disruption in this cell line. Also, what stands out from the result is that s-GO affected the migration of BEAS-2B and HeLa cells differently, with the directional persistence lowered in the former and the cell speed reduced in the latter. The result showed that the effect of s-GO on cell migration was also cell-line specific, further supporting the hypothesis that GO induces differential biological changes in cancer and non-cancer cells.

In addition, as shown in **Figure 7**, we directly compared the migratory ability between BEAS-2B, NIH/3T3, HeLa and A549 cells when interacting with s-GO (50 μg/mL). The non-cancer cells exhibited a larger spreading area compared to the cancer cell lines, with BEAS-2B cells exploring the widest territory, followed by NIH/3T3, A549 and HeLa cells (**Figure 7a**). This agrees with the MSD plot (**Figure 7b**). Furthermore, HeLa and A549 cells displayed a higher directional persistence than BEAS-2B and NIH/3T3 cells. The mean directionality ratio for HeLa and A549 cells was 0.382 ± 0.005 and 0.322 ± 0.006, whereas for BEAS-2B and NIH/3T3 cells was 0.263 ± 0.010 and 0.260 ± 0.008, respectively (**Figure 7c**). In contrast, BEAS-2B and NIH/3T3 cells displayed a higher cell speed than the A549 and HeLa cells when interacting with s-GO (**Figure 7d**). For example, the average speed for BEAS-2B and NIH/3T3 cells was 0.640 ± 0.280 and 0.327 ± 0.093 μm/min, respectively. In comparison, the speed for HeLa and A549 cells was 0.117 ± 0.068 and 0.114 ± 0.056 μm/min, respectively.

**Figure 7:**
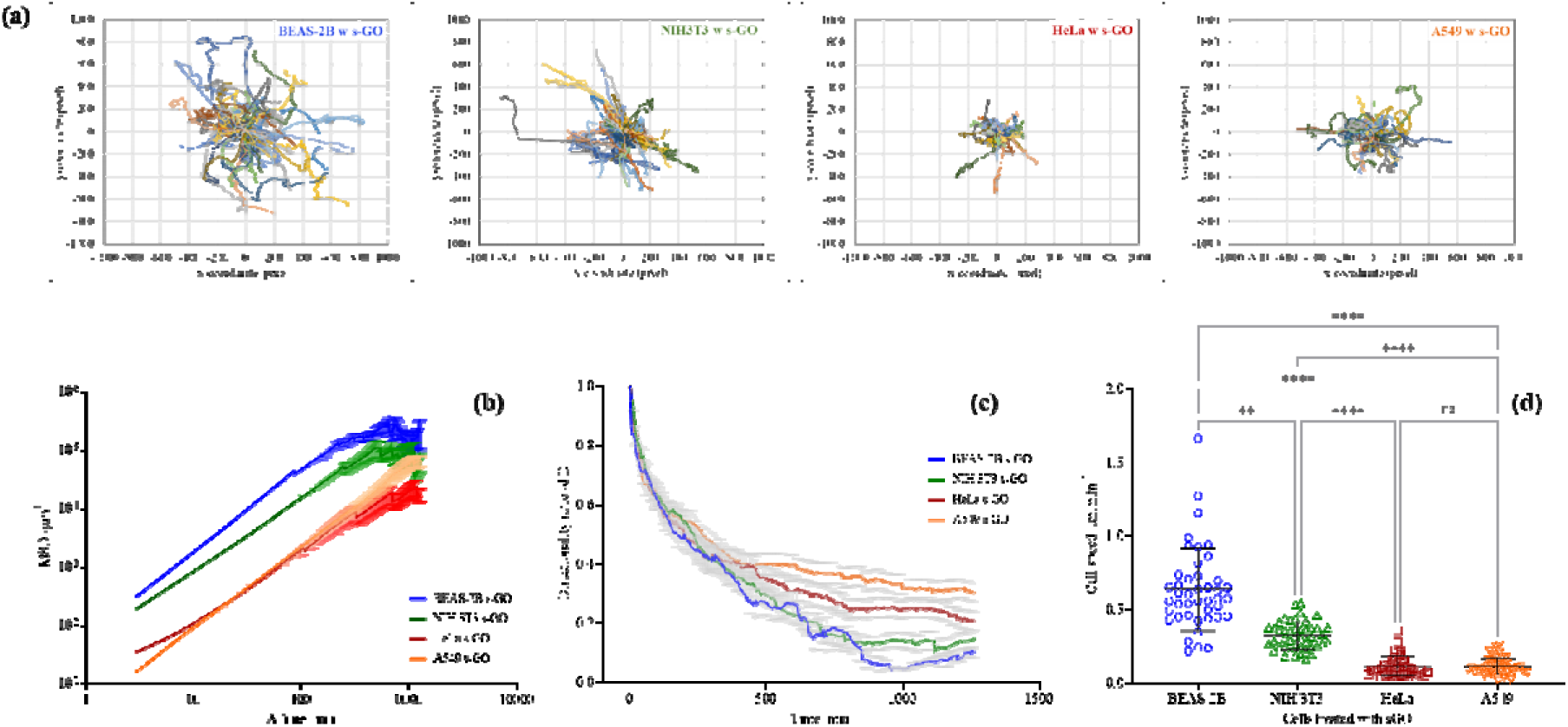
Comparison of cancer (HeLa, A549) and non-cancer cells (BEAS-2B, NIH/3T3) migration when interacting with s-GO (50 μg/mL). Cell trajectories are visualised *via* **(a)** plot at the origins and further analysed *via* **(b)** MSD, **(c)** directionality analysis, and **(d)** cell speed measurement. Cell speeds were statistically analysed using analysis of variance (one-way ANOVA) with Dunn’s multiple comparisons test. Number of cells analysed per condition ≥ 43. *Statistically different: *\*\* p* < 0.01, *\*\*\*\*p* < 0.0001, ns = not significant.

Taken together, direct comparisons of the migratory ability between BEAS-2B, NIH/3T3, HeLa and A549 cells confirmed that a greater effect of GO on cancer cell migratory ability than non-cancer cells (see **Figure 7b**). BEAS-2B and NIH/3T3 non-cancer cells followed a more curved cell path and moved faster than the HeLa and A549 cancer cells when interacting with s-GO. This result supports GO induced actin filament disruption in cancer cells and further illustrates differences between cancer and non-cancer cells when interacting with the material. Consistent with our result, studies have reported the ability of GO to retard cell migration^13,20,41,58^. However, most of these studies were performed only on cancer cells and lacked systematic comparison with non-cancer cells. Considering GO localised mostly extracellularly to cancer cells but was able to induce intracellular (see **Figure 5**, **S20** and **21**), and behaviour (see **Figure 6**) changes in cancer cells, elucidating the complex linkage between GO and cancer cells provides a novel opportunity to advance the understanding of manipulating cancer cells with extracellularly localised GO.

## Conclusion

We systematically compared interactions of non-labelled, thin GO flakes with eight cancer and five non-cancer cell lines using a mix-method approach (CLSM, TEM, flow cytometry). Large, small and ultra-small GO flakes were found to predominately interact with the plasma membrane of cancer cells, rather than being internalised. Efficient uptake of GO is reported in all studied non-cancer cell lines.

We explored factors that drive cell-type preference towards GO uptake, including material properties and the effect of GO on endocytosis-relevant biological properties of the cell. Uptake of other materials, including polystyrene microspheres and graphene flakes with various sizes and surface charges, reveals that all were internalised by cancer (HeLa, A549) and non-cancer (Beas-2B, NIH3T3) cells. This confirmed that cancer cells could take up different NMs, including thicker graphene flakes, revealing the role of GO thickness in the observed uptake differences. Furthermore, we showed that GO induced actin cytoskeleton (a key player in endocytosis) disruption in cancer but not in non-cancer cells, is leading to differential changes in the migratory behaviour of cancer cells.

Our results reveal that an interplay between properties of GO (in particular, thickness of the material) and actin cytoskeleton disruption contributed to the observed uptake differences between cancer and non-cancer cells. This work invites further studies to explore the effects of GO on plasma membrane of cancer cells, potentially translating into an opportunity towards extracellular cancer cell manipulation using GO and plasma membrane targeted biomedical applications.

## Experimental

### Materials characterisation

Aqueous suspensions of large (l-GO), small (s-GO), and ultrasmall (us-GO) GO sheets were produced in house, and fully characterised, as previously described in Rodrigues et al.^3^. GO stock suspensions prepared in water for injection and under endotoxin-free conditions, and the same batch has been already used in Loret et al.^76^. Emission spectra of GO (s- and us-GO, 2 mg/mL) were acquired using Cary Eclipse fluorescence spectrophotometer (Varian Inc., Agilent Technologies, UK) at room temperature. GO was prepared in Milli-Q water or cell culture medium (RPMI with 10 % FBS). GO prepared in cell culture medium was washed by centrifugation (30 min, 13 000 rpm, x 2, re-suspension in Milli-Q water) and re-suspended in Milli-Q water for measurement (excitation wavelength = 525 nm, blank = Milli-Q water). Graphene dispersion (Gr-TMA_3_ and Gr-PS1) were kindly provided by Professor Ciniza Casiraghi (University of Manchester). Full production and characterisation of Gr-TMA_3_ and Gr-PS1 can be found in Shin et al. and Yang et al.^30,49,50^.

### Cell culture

Thirteen immortalised cell lines were used in this study. This includes five normal cell lines and eight cancer cell lines. (1) Human epithelial bronchial cells (BEAS-2B, CRL-9609, ATCC, LGC standards, UK), (2) human prostate epithelial cells (PNT-2, kindly provided by Professor Anna Nicolas, University of Manchester), and (3) human prostate carcinoma cells (LNCaP, kindly provided by Professor Anna Nicolas, University of Manchester) were maintained in RPMI-1640 cell culture medium (R8758, Sigma-Aldrich, Merck Sigma, UK). (4) Human epithelial colon adenocarcinoma cells (SW 480, CCL-228, ATCC, LGC standards. UK) were maintained in advanced RPMI-1640 cell culture medium (R8785, Sigma-Aldrich, Merck Sigma, UK). (5) Mouse fibroblast embryonic cells (NIH/3T3, CRL-1658, ATCC, LGC standards, UK), (6) human epithelial keratinocyte cells (HaCaT, PCS-200-011, ATCC, LGC standards, UK), (7) human epithelial embryonic kidney cells (293T, CRL-2316, ATCC, LGC standards, UK), (8) human epithelial cervical adenocarcinoma cells (HeLa, CCL-2, ATCC, LGC standards, UK), and (9) human epithelial glioblastoma cells (U87 MG, HTB-14, ATCC, LGC standards, UK) were maintained in DMEM cell culture medium (D6429, Sigma-Aldrich, Merck Sigma, UK). (10) Human epithelial lung carcinoma cells (A549, CCL-185, ATCC, LGC standards, UK), and (11) human epithelial prostate adenocarcinoma cells (PC3, CRL-1435, ATCC, LGC standards, UK) were maintained in F-12 Ham cell culture medium (N6658, Sigma-Aldrich, Merck Sigma, UK). (12) Human epithelial prostate carcinoma cells (DU 145, HTB-81, ATCC, LGC standards. UK), and (13) human epithelial bone marrow neuroblastoma cells (SH-SY5Y, CRL-2266, ATCC, LGC standards, UK) were maintained in EMEM cell culture medium (M4655, Sigma-Aldrich, Merck Sigma, UK). All the cell lines except for LNCaP cells, were supplemented with 10 % FBS, 1000 units Penicillin, and 1 mg/mL Streptomycin. LNCaP cells were supplemented with 10 % hyclone FBS, 1 mM sodium pyruvate and 10 mM of HEPES solution. The cells were maintained at 37 °C in a humidified 5 % CO_2_ incubator. Cells were split at 80 % confluence with 0.05 % Trypsin-EDTA (Sigma-Aldrich, Merck Sigma, UK), and 10 % FBS (or hyclone FBS for LNCaP cells) was used to stop the activity of Trypsin-EDTA.

### Cell culture treatments

Cells were seeded in Cellview™ dishes, 12-well plates, or 24-well plates with Aclar for experiments by confocal microscopy, flow cytometry, and TEM, respectively. Cells were always seeded in the cell-type specific medium for at least 24 h before treatments and treated when the cells reached 60 – 80 % confluence (except for the wound healing assay). In all treatment condition, RPMI-1640 cell culture medium supplemented with 10 % FBS, 1000 units Penicillin, and 1 mg/mL Streptomycin were used as the pre-treatment and treatment medium and maintained at 37 °C in a humidified 5 % CO_2_ incubator.

### Confocal microscopy

#### Uptake of GO

Cells were treated with GO (l-GO at 50 µg/mL, s-GO and us-GO at 25, 50, and 75 µg/mL, 0.5 mL/well, 24 h), after removal of cell supernatant, stained with the CellMask™ green plasma membrane dye (C37608, Thermo Fisher Scientific, UK) in the fresh treatment medium with a dilution of 1:2500, and imaged by Zeiss 780 CLSM using the 40X objective. Zeiss microscope software ZEN was used to process all the images. Excitation/emission wavelength: CellMask™ green = 488/520, GO = 594/620-690 nm.

#### Uptake of the beads

Cells were treated with the positively charged (+1 µm and +0.2 µm) or negatively charged (−1 µm, −0.5 µm and −0.1 µm) beads (1.5 µL/mL, 0.5 mL/well, 24 h, F8765, F8764, F8823, F8813, F8803, Thermo Scientific, UK, respectively), after removal of cell supernatant, stained with CellMask™ deep red plasma membrane dye (C10046), Thermo Fisher Scientific, UK) in the fresh treatment medium with dilution of 1:2500, and imaged by Zeiss 780 CLSM using the 40X objective. Zeiss microscope software ZEN was used to process all the images. Excitation/emission wavelength: CellMask™ deep red = 649/666, beads = 505/515 nm.

#### Uptake of graphene flakes

Cells were treated with the positively charged (Gr-TMA_3_) or negatively charged (Gr-PS1) graphene flakes (50 µg/mL, 0.5 mL/well, 24 h), just before imaging, cell supernatant removed and stained with Fluorescein Diacetate live cell dye (100 nM, 0.5 mL/well, Thermo Fisher Scientific, UK) in the fresh treatment medium, and imaged by Zeiss 780 CLSM using the 40X objective. Zeiss microscope software ZEN was used to process all the images. Excitation/emission wavelength: FDA = 488/520 nm.

#### Time-lapse tracking: untreated vs s-GO treated cells

BEAS-2B or HeLa cells were seeded in the Cellview™ dish and treated with s-GO (50 μg/mL, 0.5 mL/well) containing CellMask™ green plasma membrane stain (C37608, Thermo Scientific, UK). Live-cell time lapse was recorded within 1 h after the addition of GO treatment for 21 h, using Zeiss 780 CLSM (40X objective, time lapse mode). The videos were then extracted using Zeiss microscope software ZEN with the compression ratio of 20 and frames per second of 10. Excitation/emission wavelength: CellMask™ green = 488/520, GO = 594/620-690 nm.

#### Time-lapse tracking: cancer vs non-cancer cells when interacting with s-GO

BEAS-2B, HeLa and A549 cells were seeded in the same Cellview™ dish and treated with s-GO (50 μg/mL, 0.5 mL/well) containing CellMask™ green plasma membrane stain (C37608, Thermo Scientific, UK). Live-cell time lapse was recorded within 1 h after the addition of GO treatment for 21 h, using Zeiss 780 multiphoton CLSM (40X objective, time lapse mode). The videos were then extracted using Zeiss microscope software ZEN. Excitation/emission wavelength: CellMask™ green = 488/520, GO = 594/620-690 nm.

#### Time-lapse tracking: video processing and analysis

To obtain the cell trajectory information, cells were tracked manually using the CellTracker (version 1.1) software^77^. Briefly, the extracted video of live-cell time lapse by Zeiss 780 multiphoton CLSM was loaded onto CellTracker as Bio-Format images. Then, using the manual tracking option, the position of individual cell was manually defined by user clicking on the cell at every image frame until cell leave the frame; ≥ 43 cells were analysed in every conditions. Afterward, the statistic tap in CellTracker was selected, user defined manually the pixel size of frame (0.209 μm/pixel) and the time elapsed between subsequent frames (2.88 min/frame) according to the pre-defined setting of the recorded time lapse. The average cell speeds (μm/min) and individual cell trajectory (defined as *xy*-coordinate in each image frame) were extracted using the statistic function from CellTracker. Subsequently, the computed *xy*-coordinates data from the CellTracker were further analysed using the DiPer programme^75^ to generate the plot at the origin, MSD curve, and directionality curve. Briefly, the pre-written program codes for DiPer were downloaded and saved as Maro-Enabled Workbook using Excel^75^. Then the *xy*-coordinates data from the CellTracker were copied onto the Maro-Enabled Workbook and further analysed using the corresponding Macros for the plot at origin, MSD curve, and directionality curve^75^.

#### Staining of the actin filament

Cells were treated with GO (s-GO and us-GO, 100 μg/mL, 0.5 mL/well, 24 h) or M /JCD (5 mM, 0.5 mL/well, 4 h, C4555, Sigma), washed with pre-warmed PBS (0.5 mL/well, D8662, Sigma-Aldrich, Merck Sigma, UK), fixed with formaldehyde (3.7 %, 0.5 ml/well, 10 min; 28908 Thermo Fisher Scientific, UK), permeabilized with Triton-X (0.1 % in PBS (−/−), 0.5 mL/well, 10 mins), re-washed with PBS (−/−) (0.5 mL/well, x 2, D8537, Sigma-Aldrich, Merck Sigma, UK), stained with Alexa Fluor™ 488 Phalloidin (1:1500 dilution prepared in PBS (−/−), 0.5 mL/well, 20 min, A12379, Thermo Fischer, UK). After staining, cells were washed with PBS (−/−) (0.5 mL/well, x 2) and ProLong™ Gold Antifade Mountant (P36930, Thermo Fisher Scientific, UK) was added to the cells. The cells were exposed to a higher concentration of GO (100 μg/mL) because the potential role of GO as an actin filament disrupting agent was investigated. Following the working principle for pharmaceutical inhibitors, the highest tested non-toxic concentration of inhibitor was used^28^.

Cells were imaged using Zeiss 780 CLSM with the 40X objective. Images were processed using Zeiss microscope software ZEN. Excitation/emission wavelength: Phalloidin = 495/518.

### Flow cytometry

#### Cellular interactions with GO

Cells were treated with s-GO or us-GO (50 µg/mL, 1 mL/well, 24 h). After treatment, cells were collected with and without washing. The cells without washing were detached with 0.05 % Trypsin-EDTA (300 uL/well, 10 min), neutralised with FBS (30 uL/well), collected in 1.5 mL Eppendorf tube, and stored in ice until analysis. The cells with washing were washed with PBS (−/−) (1 mL/well, x 2), detached with 0.05 % Trypsin-EDTA (300 uL/well, 10 min), neutralised with FBS (30 uL/well), collected in 1.5 mL Eppendorf tube, re-washed by centrifugation (500 RPM, 5 mins, 4 ^O^C, x 2), re-suspended in PBS (−/−) (300 uL/tube), and stored in ice until analysis. The samples were sampled using FACSVerse flow cytometry using PE-Cy7-A channel (band pass: 488 780/60). Excitation/emission band pass: GO = 594/620-690.

#### Cellular interactions with beads

Cells were treated with the positively charged (+1 µm and +0.2 µm) or negatively charged (−1 µm, −0.5 µm and −0.1 µm) beads (1.5 µL/mL, 1 mL/well, 24 h, F8765, F8764, F8823, F8813, F8803, Thermo Scientific, UK, respectively). After treatment, detached with 0.05 % Trypsin-EDTA (300 uL/well, 10 min), neutralised with FBS (30 uL/well), collected in 1.5 mL Eppendorf tube, stored in ice, analysed by FACSVerse flow cytometry (bandpass: 488 530/30) using the FITC channel. Excitation/emission band pass: +1, +0.2, −1, −0.5, −0.1 µm beads= 505/515.

### TEM

Cells were seeded on sterilised Aclar placed in 24-well plate. Subsequently, cells were treated with s-GO or us-GO (50 μg/mL, 0.5 mL/well, 24 h) and fixed at room temperature (4 % formaldehyde, 2.5 % glutaraldehyde in 0.1 M HEPS budder, 0.5 mL/well, 1 h). Samples were postfixed with 1 % osmium tetroxide + 1.5 % potassium ferrocynaide in 0.1M cacodylate buffer (pH 7.2) for 1 hour, then in 1 % uranyl acetate in water overnight. Then they were dehydrated in ethanol series infiltrated with TAAB LV resin and polymerised for 24h at 60C degree. Sections were cut with Reichert Ultracut ultramicrotome and observed with FEI Tecnai 12 Biotwin microscope at 100kV accelerating voltage. Images were taken with Gatan Orius SC1000 CCD camera.

### Statistical analysis

Flow cytometry, cell speed, and directionality data were statistically analysed using GraphPad Prism (version 9). The analysis of variance (one-way (**Figure 4**) or two-way (**Figures 2e**, **S17**) ANOVA) with Tukey’s multiple comparisons test was used for flow cytometry data (n = 3 with duplicates). Cell speeds data were statistically analysed using unpaired T-test (**Figure 6**) or analysis of variance (one-way ANOVA) with Dunn’s multiple comparisons test and descriptive statistics (**Figure 7**), and the result was reported as mean ± SD (number of cells analysed per condition ≥ 43). Directionality data were analysed using descriptive statistics, and the result was reported as mean ± SEM. *Statistically different: * *p<*0.05, ** *p <*0.01, *** *p <*0.001 and \*\*\*\**p* < 0.0001. ns = not significant.

## Acknowledgments

We thank Dr Adrian Esteban Arranz for contributing to the synthesis of the specific GO batch used in the present study and Dr Miguel Arellano and Ms Angeliki Karakasidi for contributing to the characterization of the GO materials. The ICN2 is funded by the CERCA programme / Generalitat de Catalunya and has been supported by the Severo Ochoa Centres of Excellence programme [SEV-2017-0706] and is currently supported by the Severo Ochoa Centres of Excellence programme, Grant CEX2021-001214-S, both funded by MCIN/AEI/10.13039.501100011033. Y.C. acknowledge the studentship from the UK Research and Innovation (UKRI) Engineering and Physical Sciences Research Council (EPSRC) Centre for Doctoral Training programme (Graphene NOWNANO CDT; EP/L01548X/1). The authors acknowledge the Manchester Collaborative Centre for Inflammation Research (MCCIR) as the funding source for the FACSVerse.

## Supporting information

**Table S1:**
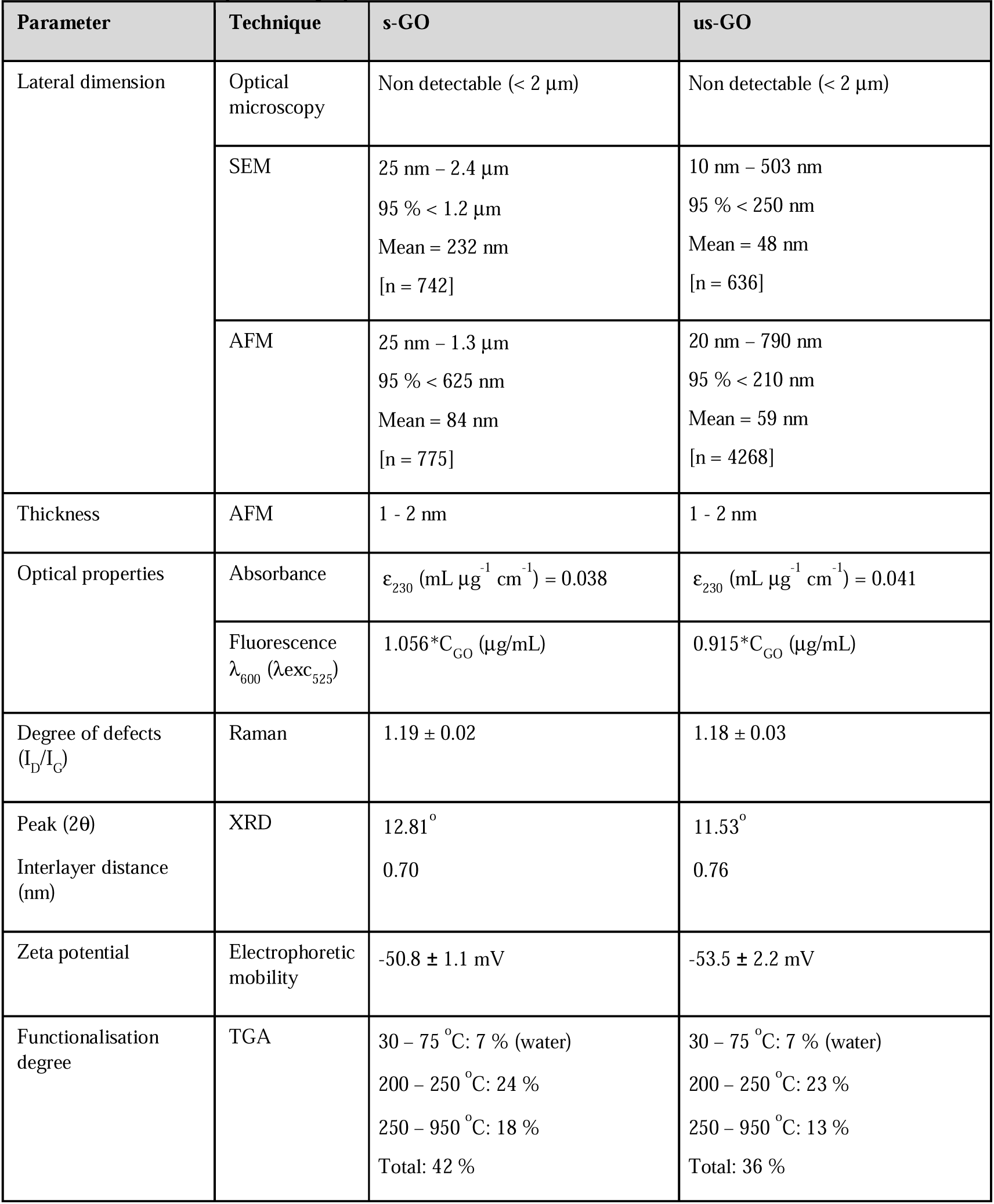
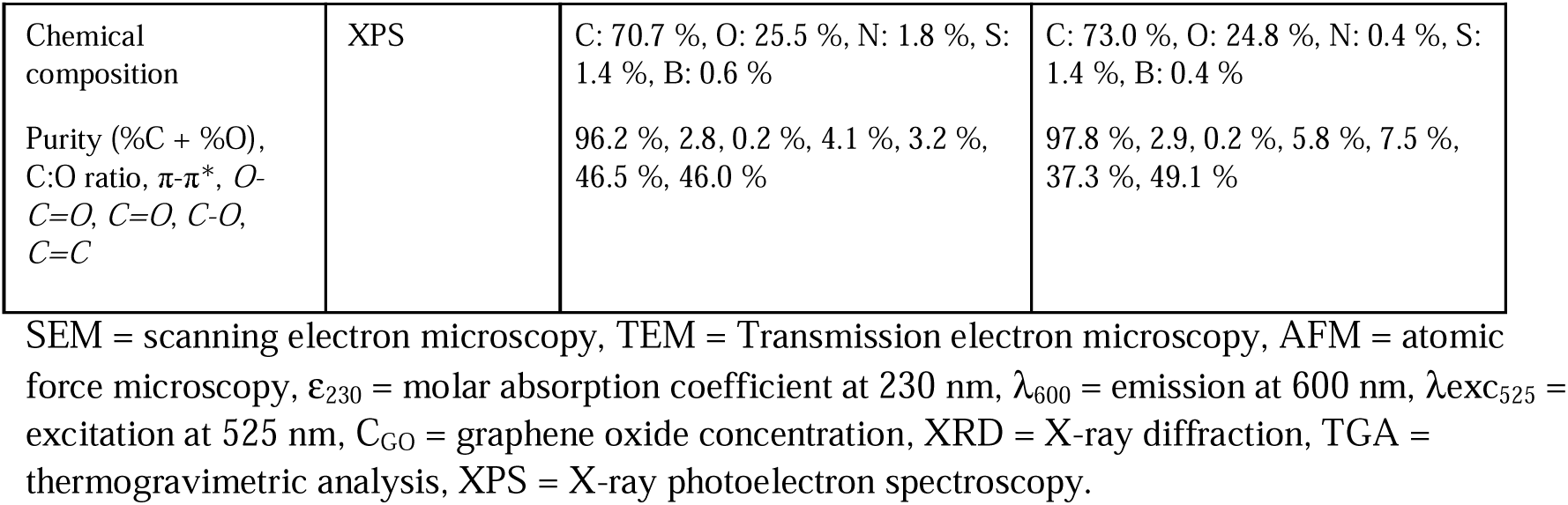
Summary of the physico-chemical characterisation of s-GO and us-GO.

| Parameter | Technique | s-GO | us-GO |
| --- | --- | --- | --- |
| Lateral dimension | Optical microscopy | Non detectable (< 2 $\mu\text{m}$) | Non detectable (< 2 $\mu\text{m}$) |
| | SEM | 25 nm – 2.4 $\mu\text{m}$<br>95 % < 1.2 $\mu\text{m}$<br>Mean = 232 nm<br>[n = 742] | 10 nm – 503 nm<br>95 % < 250 nm<br>Mean = 48 nm<br>[n = 636] |
| | AFM | 25 nm – 1.3 $\mu\text{m}$<br>95 % < 625 nm<br>Mean = 84 nm<br>[n = 775] | 20 nm – 790 nm<br>95 % < 210 nm<br>Mean = 59 nm<br>[n = 4268] |
| Thickness | AFM | 1 - 2 nm | 1 - 2 nm |
| Optical properties | Absorbance | $\epsilon_{230} \text{ (mL } \mu\text{g}^{-1} \text{ cm}^{-1}) = 0.038$ | $\epsilon_{230} \text{ (mL } \mu\text{g}^{-1} \text{ cm}^{-1}) = 0.041$ |
| | Fluorescence<br>$\lambda_{600} (\lambda_{exc} 525)$ | $1.056 * C_{GO} \text{ (}\mu\text{g/mL)}$ | $0.915 * C_{GO} \text{ (}\mu\text{g/mL)}$ |
| Degree of defects<br>($I_D/I_G$) | Raman | $1.19 \pm 0.02$ | $1.18 \pm 0.03$ |
| Peak ($2\theta$) | XRD | $12.81^\circ$ | $11.53^\circ$ |
| Interlayer distance<br>(nm) |  | 0.70 | 0.76 |
| Zeta potential | Electrophoretic mobility | $-50.8 \pm 1.1 \text{ mV}$ | $-53.5 \pm 2.2 \text{ mV}$ |
| Functionalisation degree | TGA | 30 – 75 $^\circ\text{C}$: 7 % (water)<br>200 – 250 $^\circ\text{C}$: 24 %<br>250 – 950 $^\circ\text{C}$: 18 %<br>Total: 42 % | 30 – 75 $^\circ\text{C}$: 7 % (water)<br>200 – 250 $^\circ\text{C}$: 23 %<br>250 – 950 $^\circ\text{C}$: 13 %<br>Total: 36 % |
| Chemical composition | XPS | C: 70.7 %, O: 25.5 %, N: 1.8 %, S: 1.4 %, B: 0.6 % | C: 73.0 %, O: 24.8 %, N: 0.4 %, S: 1.4 %, B: 0.4 % |
| Purity (%C + %O), C:O ratio, π-$\pi^*$, *O*-C=O, C=O, C-O, C=C | | 96.2 %, 2.8, 0.2 %, 4.1 %, 3.2 %, 46.5 %, 46.0 % | 97.8 %, 2.9, 0.2 %, 5.8 %, 7.5 %, 37.3 %, 49.1 % |

SEM = scanning electron microscopy, TEM = Transmission electron microscopy, AFM = atomic force microscopy, $\epsilon_{230}$ = molar absorption coefficient at 230 nm, $\lambda_{600}$ = emission at 600 nm, $\lambda_{exc525}$ = excitation at 525 nm, $C_{GO}$ = graphene oxide concentration, XRD = X-ray diffraction, TGA = thermogravimetric analysis, XPS = X-ray photoelectron spectroscopy.

**Figure S1:**
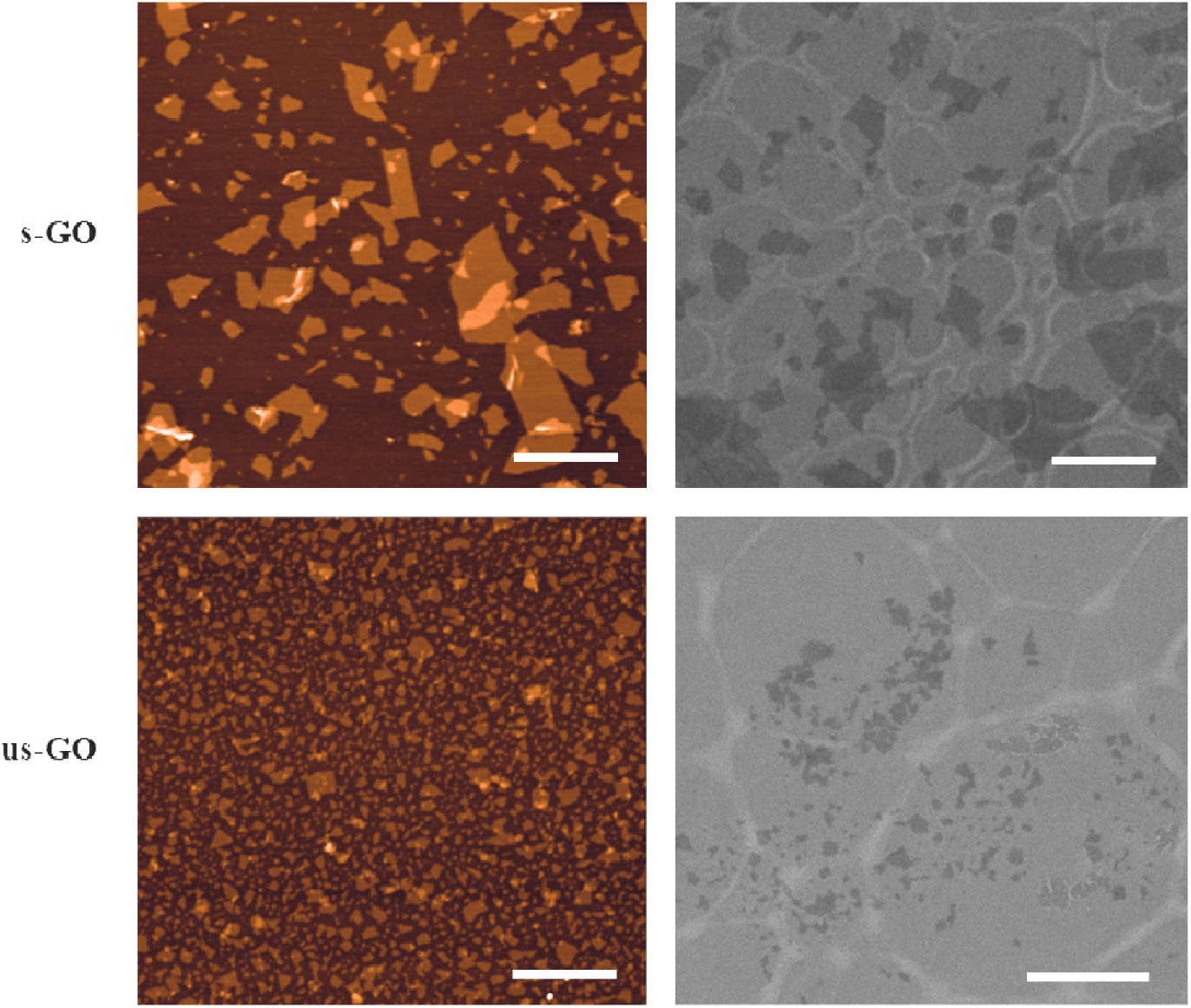
Characterisation of GO (s- and us-GO) by atomic force microscopy (left panels) and electron microscopy (right panels). Scale bar = 1 µm. See **Table S1** for the summary of the physico-chemical properties.

**Figure S2:**
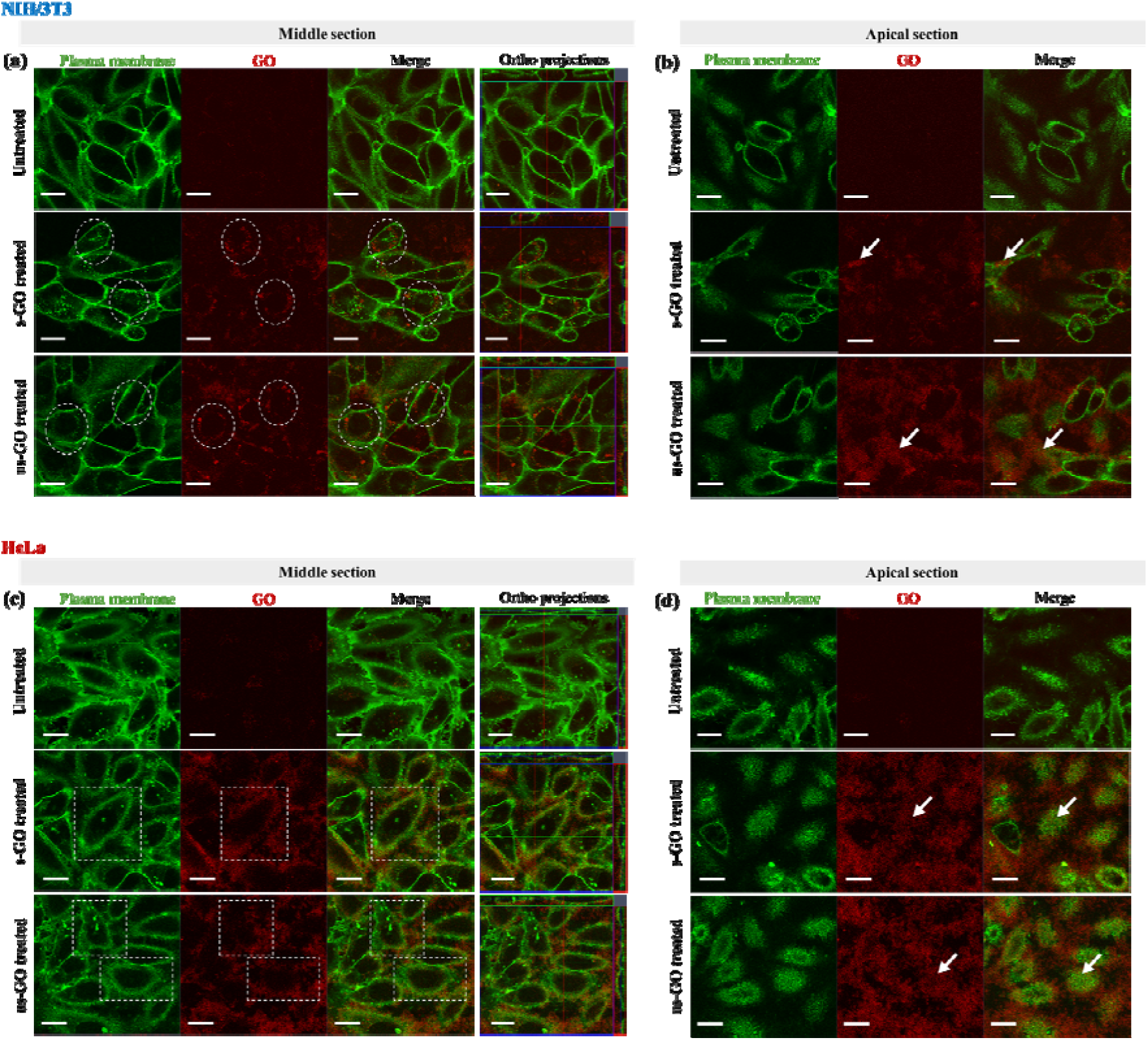
Representative images (middle and apical section) of GO (s- and us-GO) interaction profiles in **(a** – **b)** NIH/3T3 and **(c** – **d)** HeLa cells by CLSM. Both s-GO and us-GO (50 μg/mL) were found internalised in NIH/3T3 cells (indicated by white circles) but predominately surrounded the plasma membrane in HeLa cells (indicated by white rectangles). The materials were also found on the plasma membrane (indicated by white arrows) in both cell lines. Green = plasma membrane, red = GO. Scale bar = 20 µm. See **Figures S3** and **S7** for the interaction of s-GO and us-GO at different concentrations with NIH/3T3 and HeLa cells, respectively.

**Table S2:** Summary of GO (s- and us-GO) interaction profile in cancer (HeLa, A549, SW 480, SH-SY5Y, U87 MG, DU-145, PC3 and LNCaP) and non-cancer (BEAS-2B, NIH/3T3, HaCaT, 293T and PNT-2) cells by CLSM.

| Cell line | Tissue/morphology | s-GO and us-GO uptake |
| --- | --- | --- |
| **BEAS-2B** | Normal human lung/epithelial | + |
| **NIH/3T3** | Normal mouse embryonic/fibroblasts | + |
| **HaCaT** | Normal human skin/epithelial | + |
| **293T** | Normal human embryonic kidney/epithelial | + |
| **PNT-2** | Normal human prostate/epithelial | + |
| **HeLa** | Adenocarcinoma human cervix/epithelial | – |
| **A549** | Carcinoma human lung/epithelial | – |
| **SW 480** | Adenocarcinoma human cervix/epithelial | – |
| **SH-SY5Y** | Neuroblastoma human bone marrow<br>neuroblastoma/epithelial | – |
| **U87 MG** | Glioblastoma human brain/epithelial | – |
| **DU-145** | Carcinoma human prostate/epithelial | – |
| **PC3** | Adenocarcinoma human prostate/epithelial | – |
| **LNCaP** | Carcinoma human prostate/epithelial | – |

**Figure S3:**
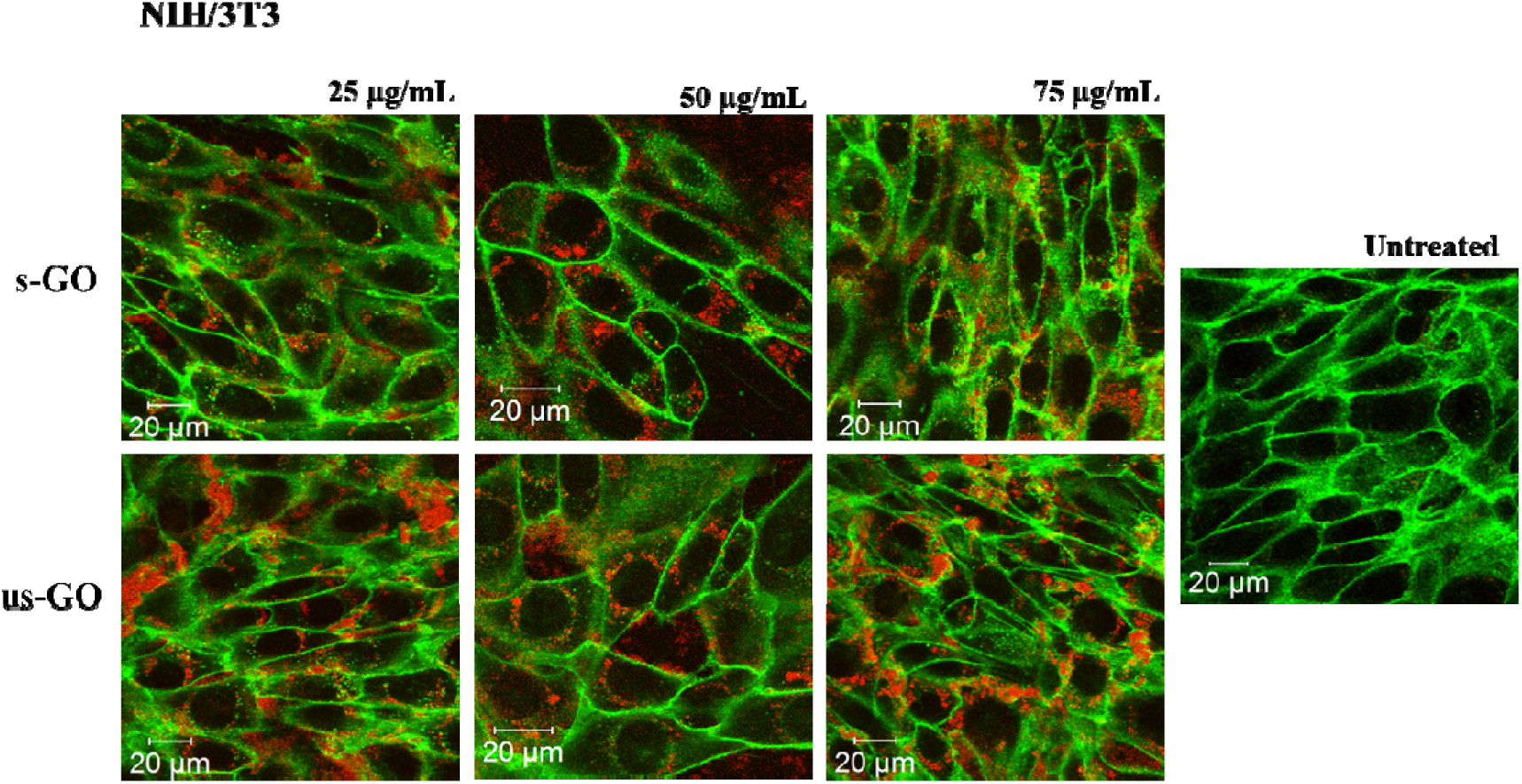
Interactions of s-GO and us-GO at 25, 50 and 75 μg/mL with NIH/3T3 cells. Green = plasma membrane, red = GO.

**Figure S4:**
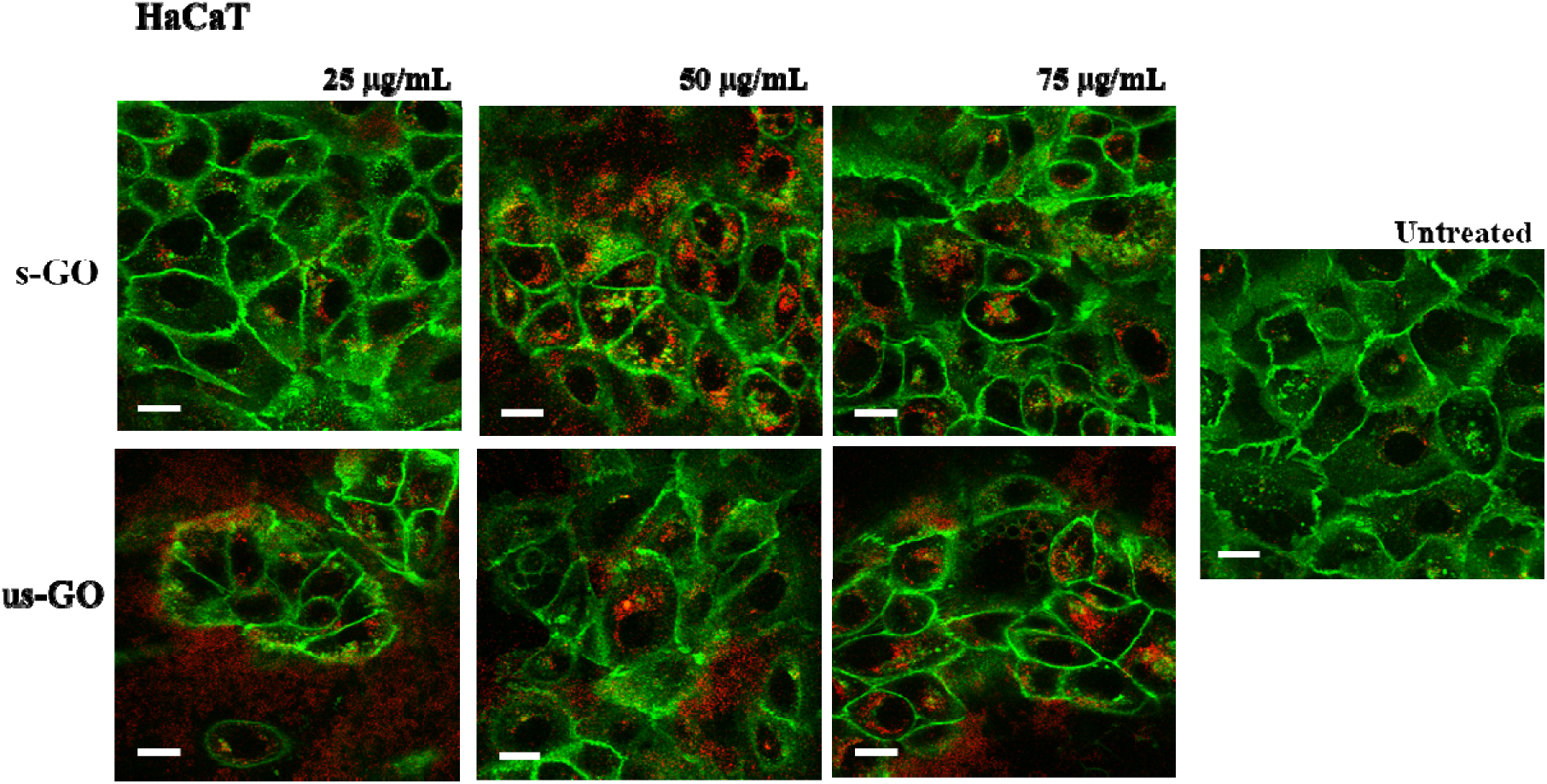
Interactions of s-GO and us-GO at 25, 50 and 75 μg/mL with HaCaT cells. Green = plasma membrane, red = GO. Scale bar = 20 μm.

**Figure S5:**
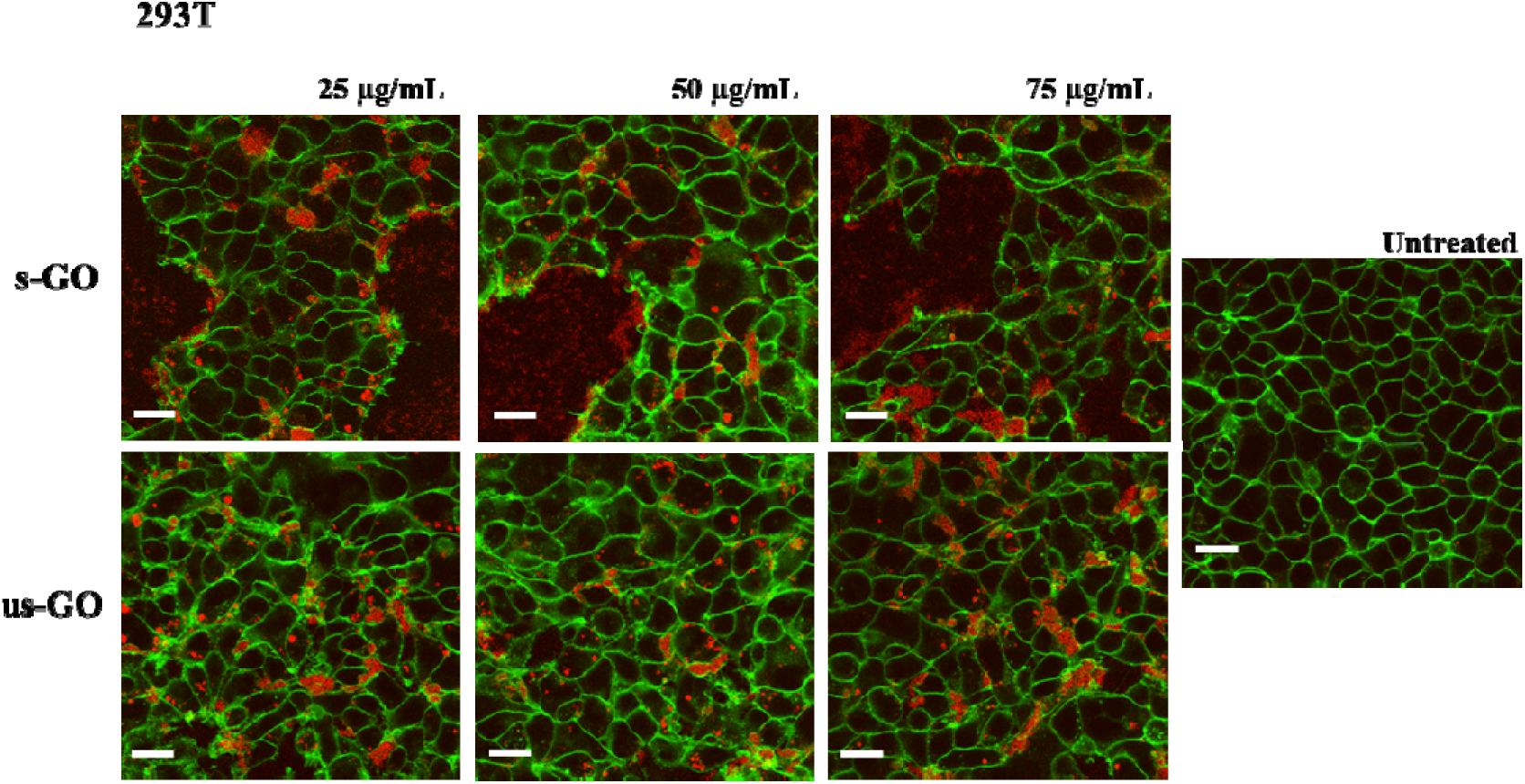
Interactions of s-GO and us-GO at 25, 50 and 75 μg/mL with 293T cells. Green = plasma membrane, red = GO. Scale bar = 20 μm.

**Figure S6:**
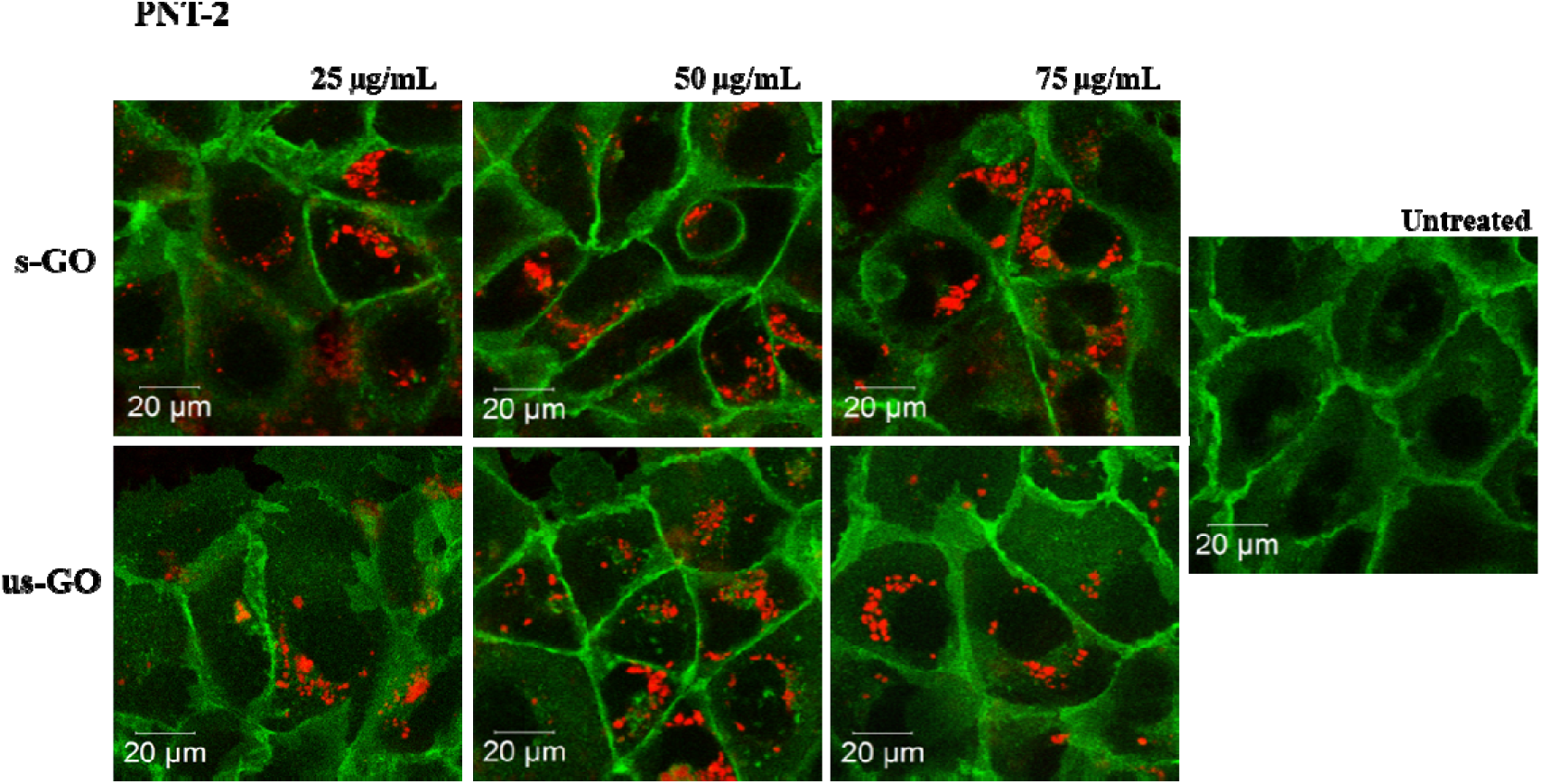
Interactions of s-GO and us-GO at 25, 50 and 75 μg/mL with PNT-2 cells. Green = plasma membrane, red = GO.

**Figure S7:**
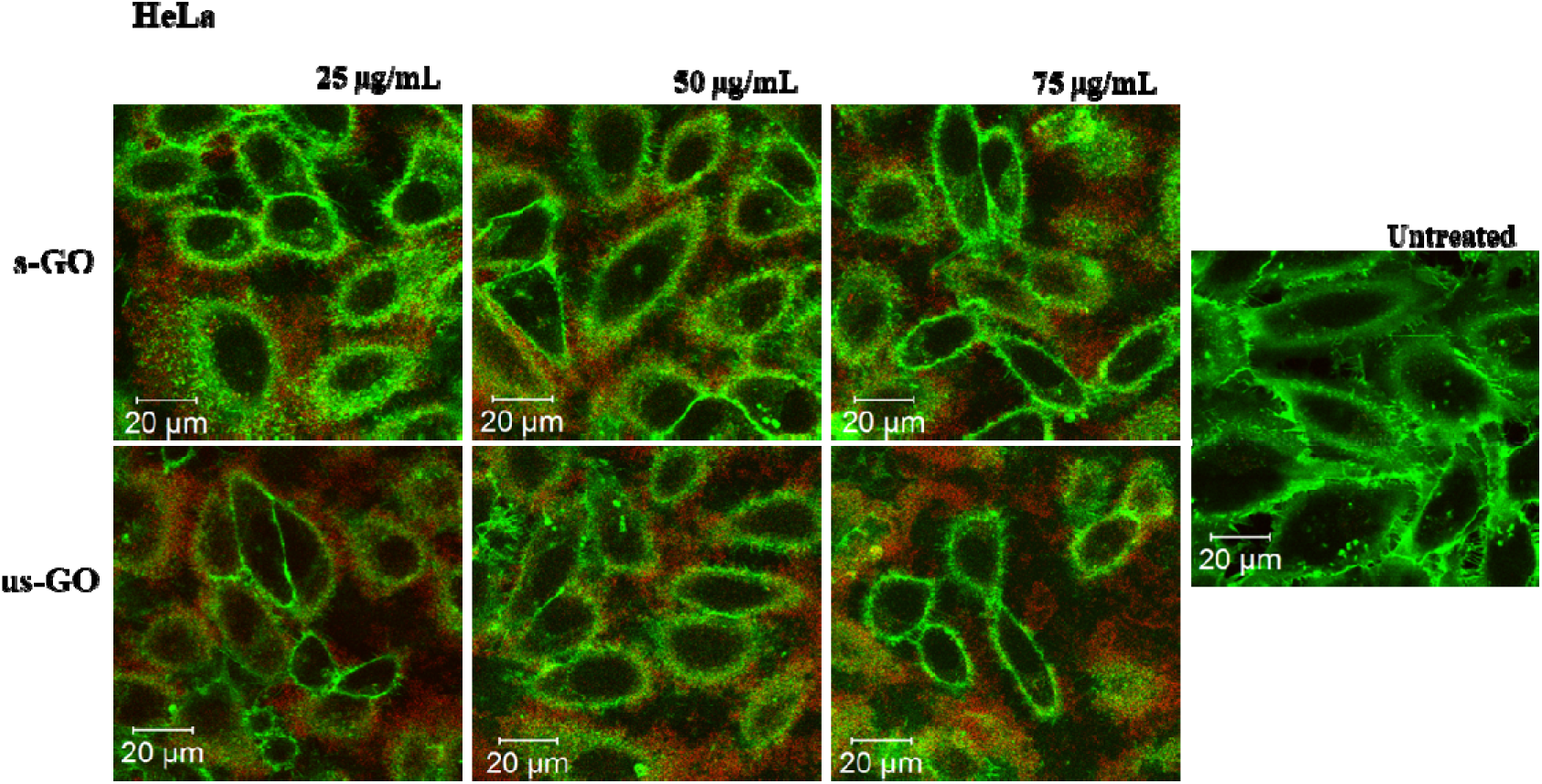
Interactions of s-GO and us-GO at 25, 50 and 75 μg/mL with HeLa cells. Green = plasma membrane, red = GO.

**Figure S8:**
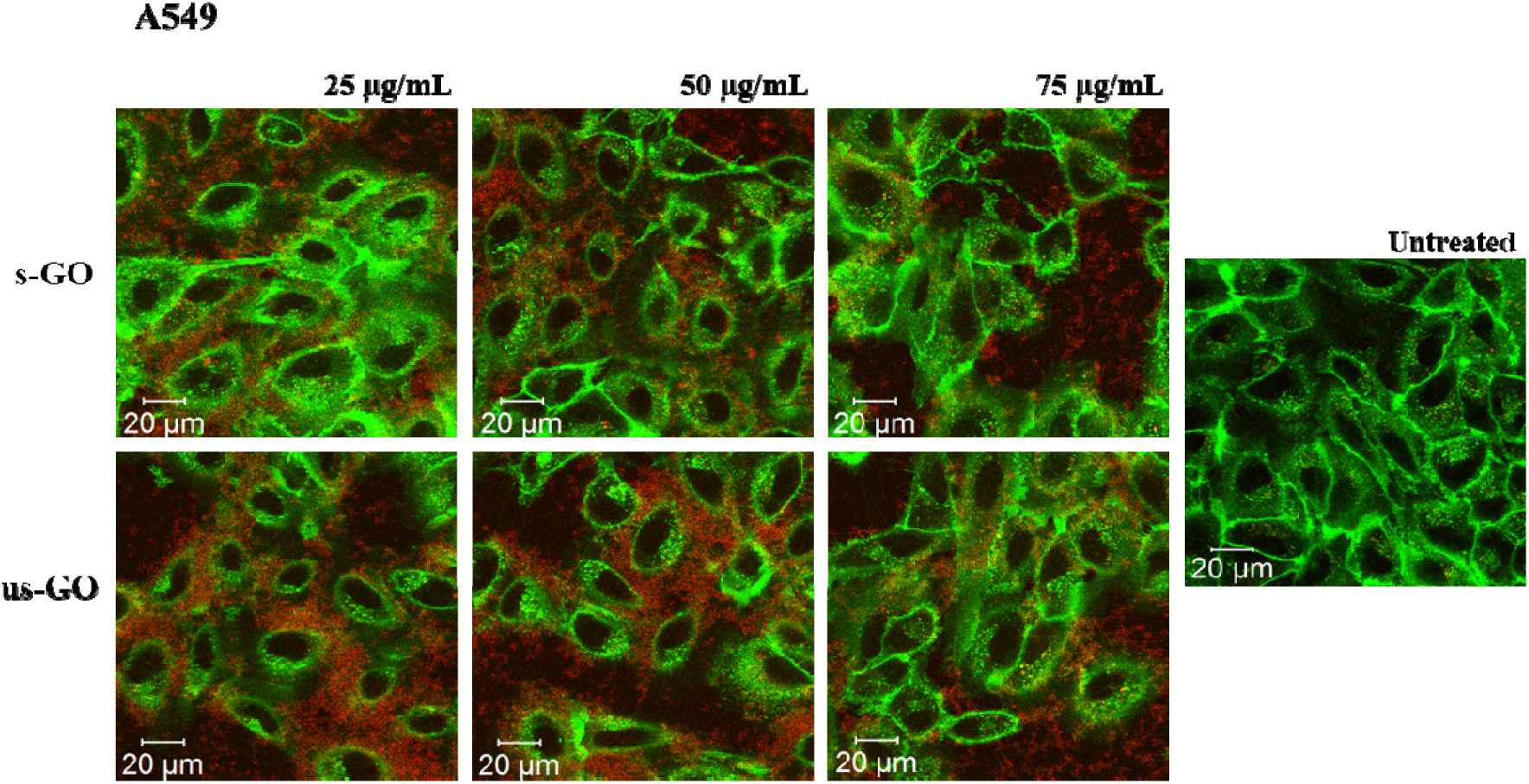
Interactions of s-GO and us-GO at 25, 50 and 75 μg/mL with A549 cells. Green = plasma membrane, red = GO.

**Figure S9:**
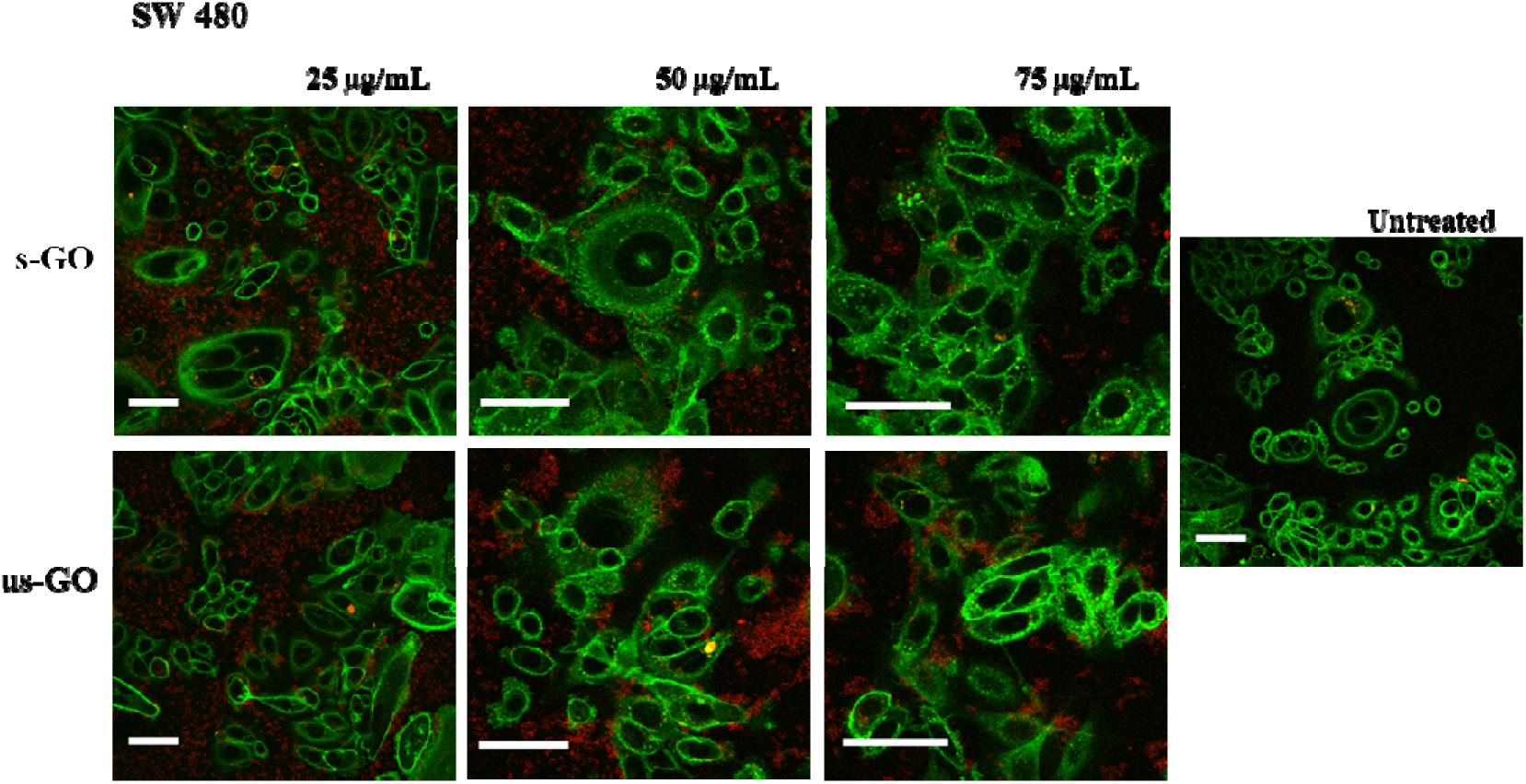
Interactions of s-GO and us-GO at 25, 50 and 75 μg/mL with SW 480 cells. Green = plasma membrane, red = GO. Scale bar = 50 μm.

**Figure S10:**
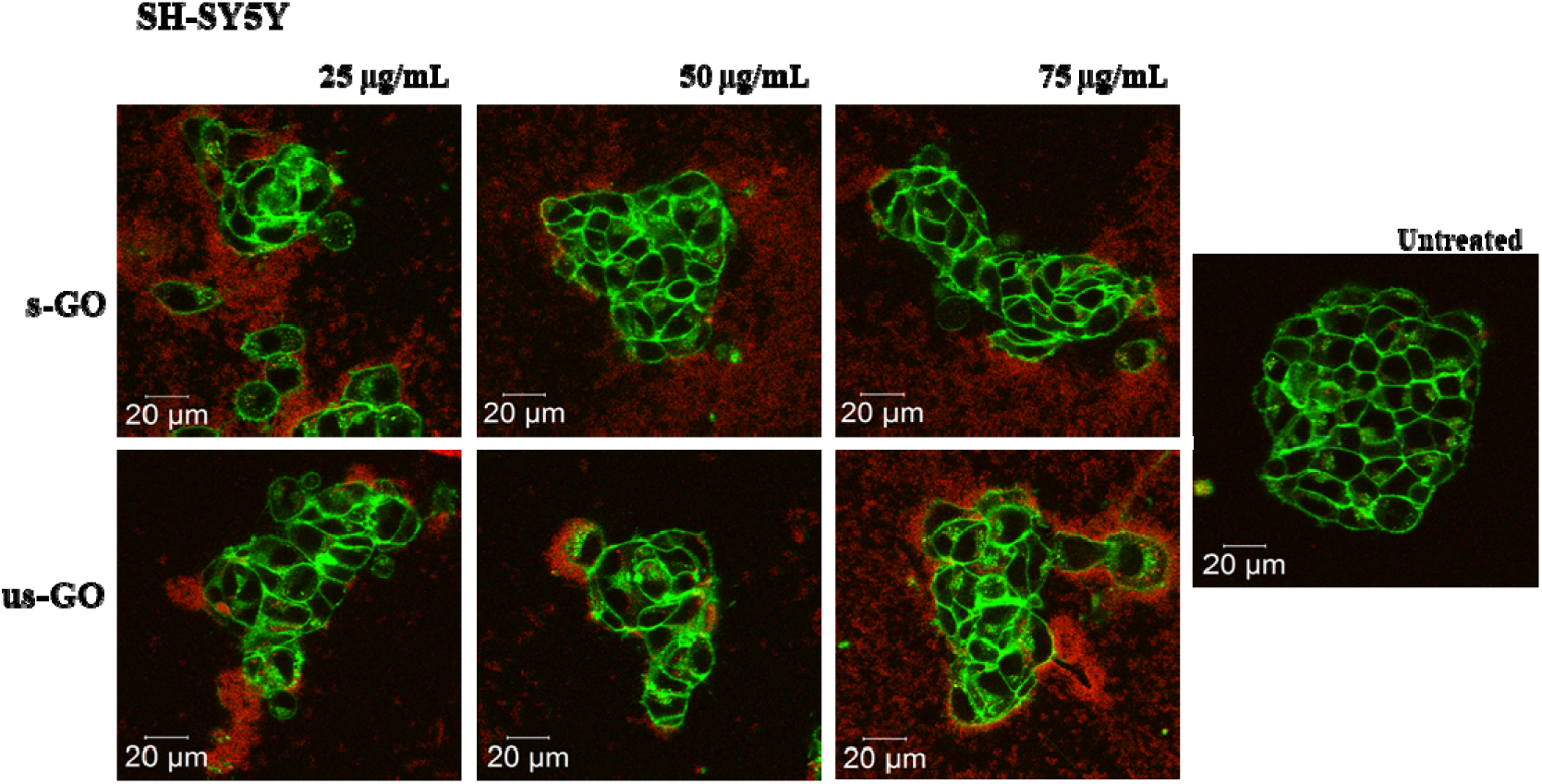
Interactions of s-GO and us-GO at 25, 50 and 75 μg/mL with SH-SY5Y cells. Green = plasma membrane, red = GO.

**Figure S11:**
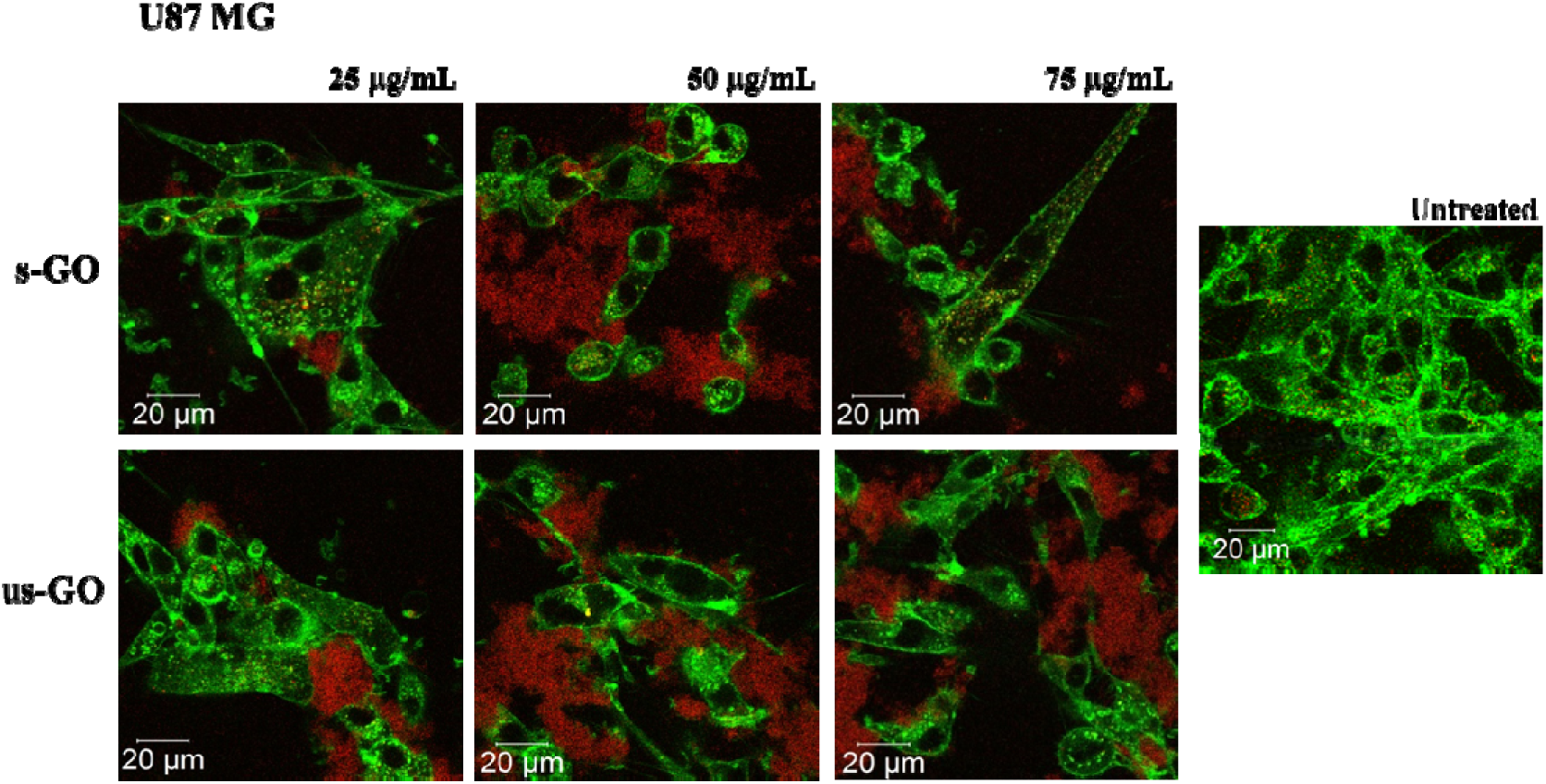
Interactions of s-GO and us-GO at 25, 50 and 75 μg/mL with U87 MG cells. Green = plasma membrane, red = GO.

**Figure S12:**
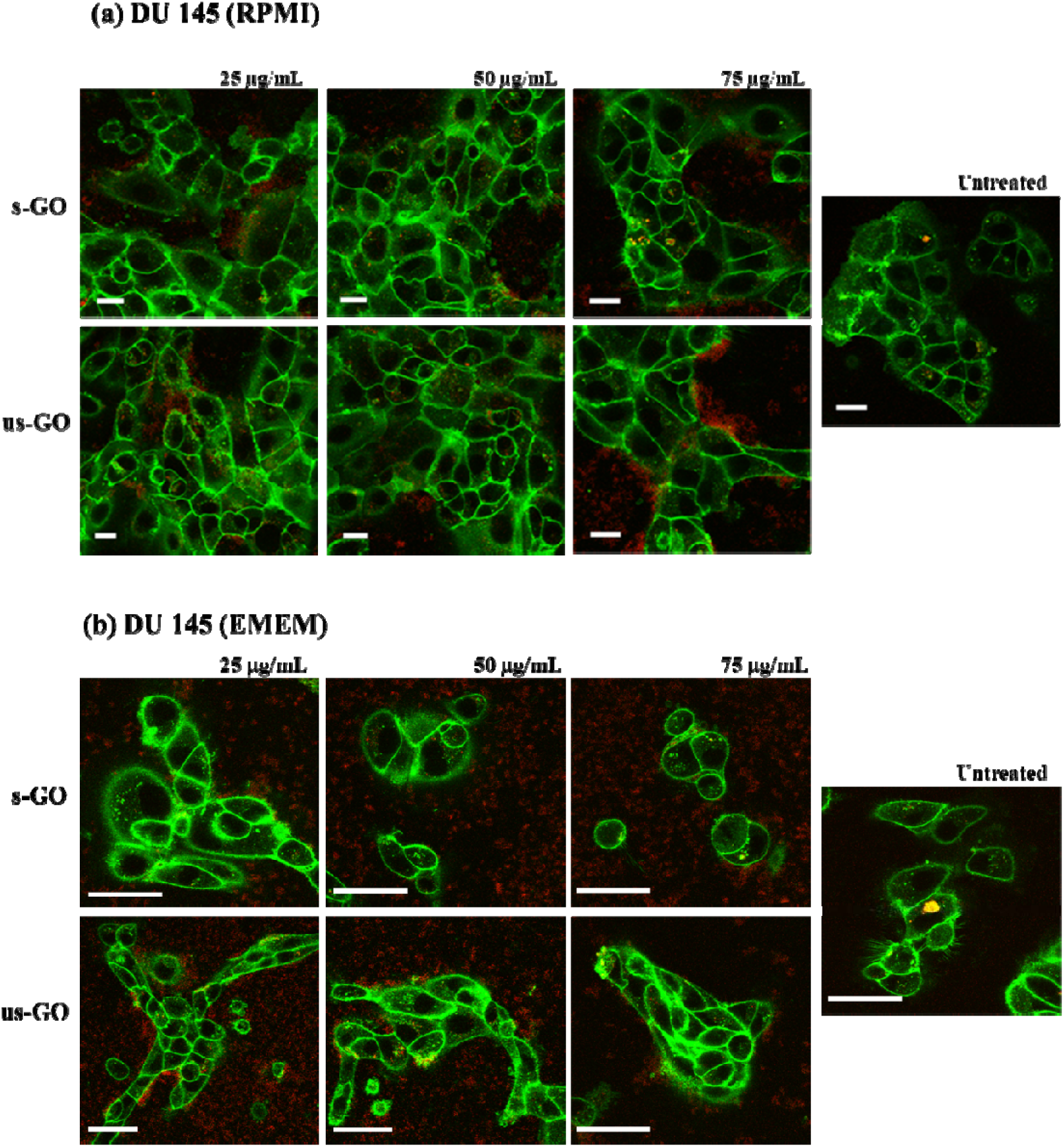
Interactions of s-GO and us-GO at 25, 50 and 75 μg/mL with DU 145 cells [in **(a)** RPMI with 10% FBS and **(b)** EMEM with 10 % FBS]. Green = plasma membrane, Red = GO. Scale bar = **(a)** 20 μm or **(b)** 50 μm.

**Figure S13:**
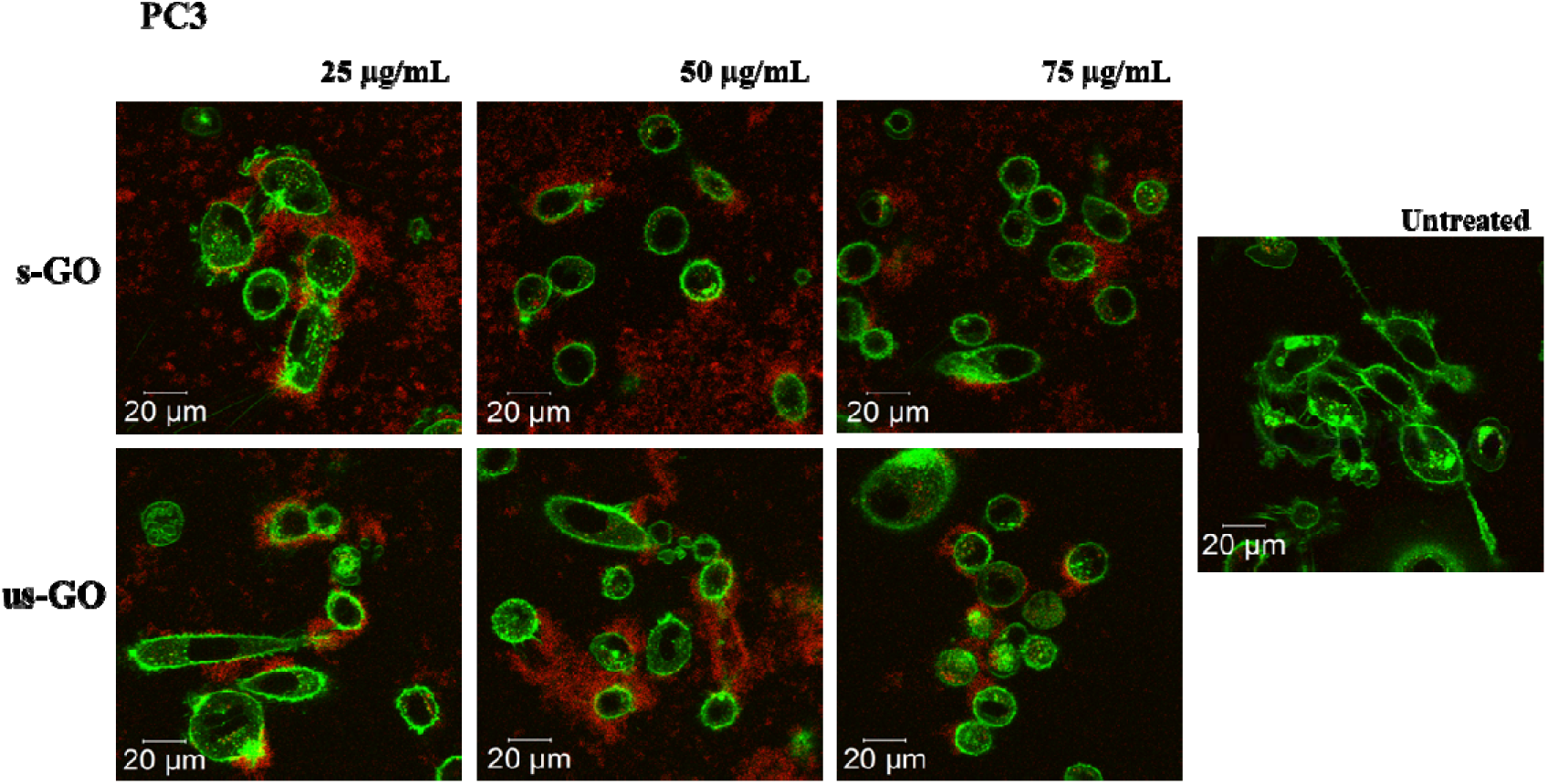
Interactions of s-GO and us-GO at 25, 50 and 75 μg/mL with PC3 cells. Green = plasma membrane, red = GO.

**Figure S14:**
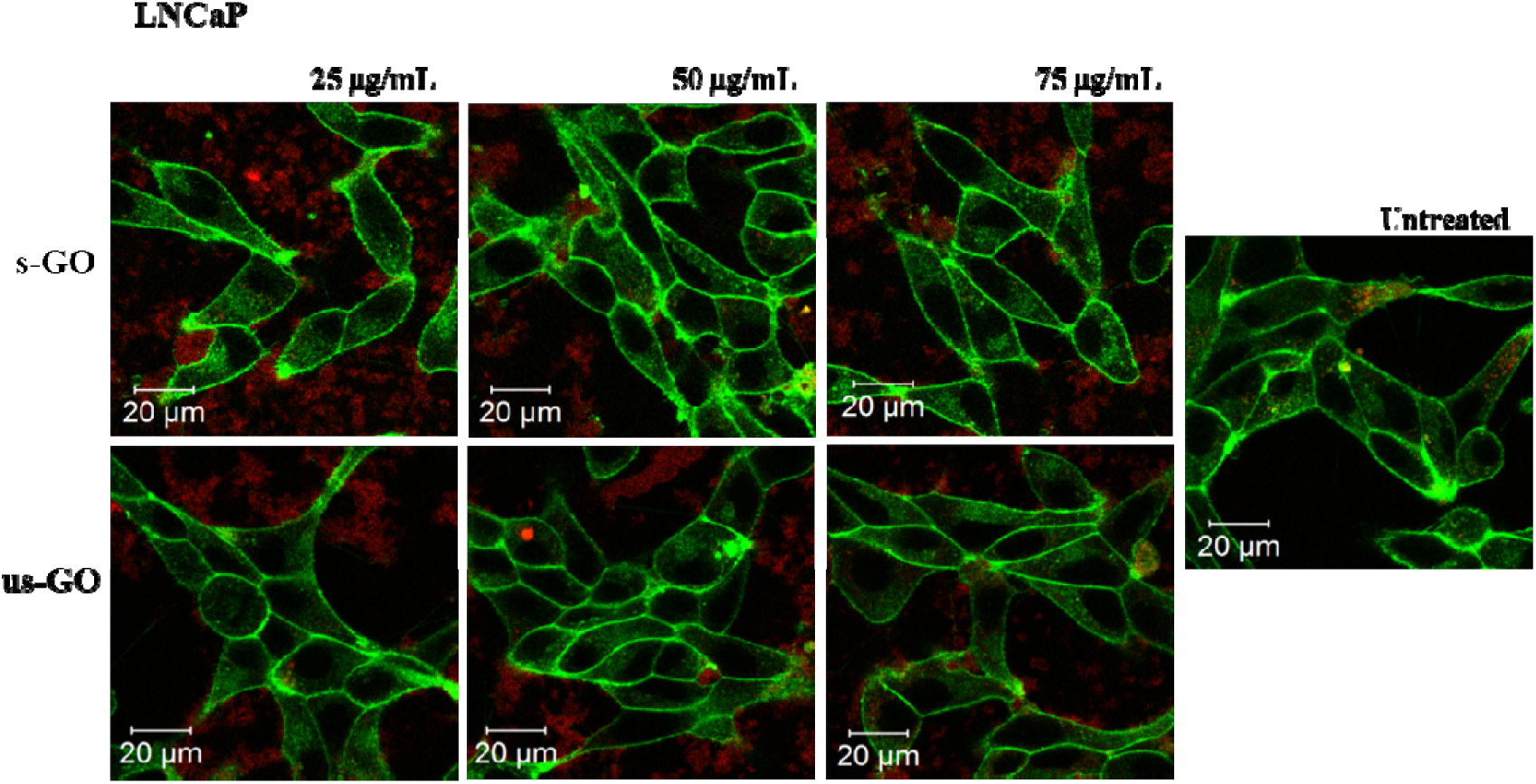
Interactions of s-GO and us-GO at 25, 50 and 75 μg/mL with LNCaP cells. Green = plasma membrane, red = GO.

**Figure S15:**
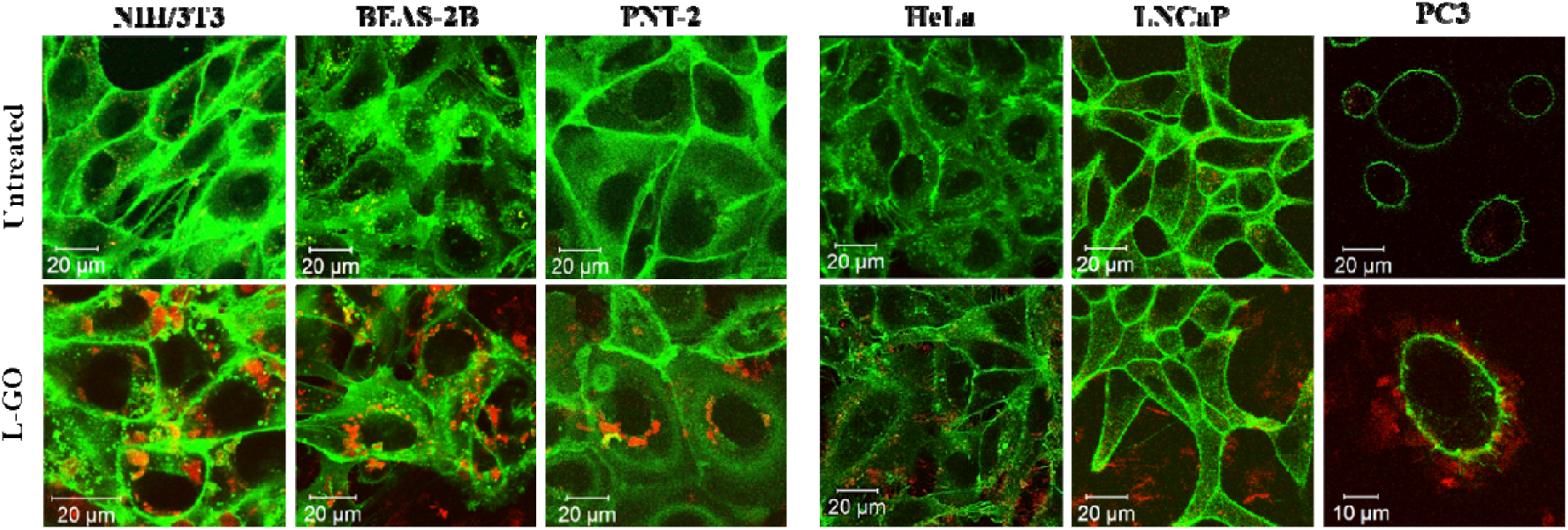
Interaction l-GO (50 μg/mL) with non-cancer (NIH/3T3, BEAS2B, PNT-2) and cancer (HeLa, LNCaP and PC3) cell lines by confocal imaging. Green = plasma membrane, red = GO.

**Figure S16:**
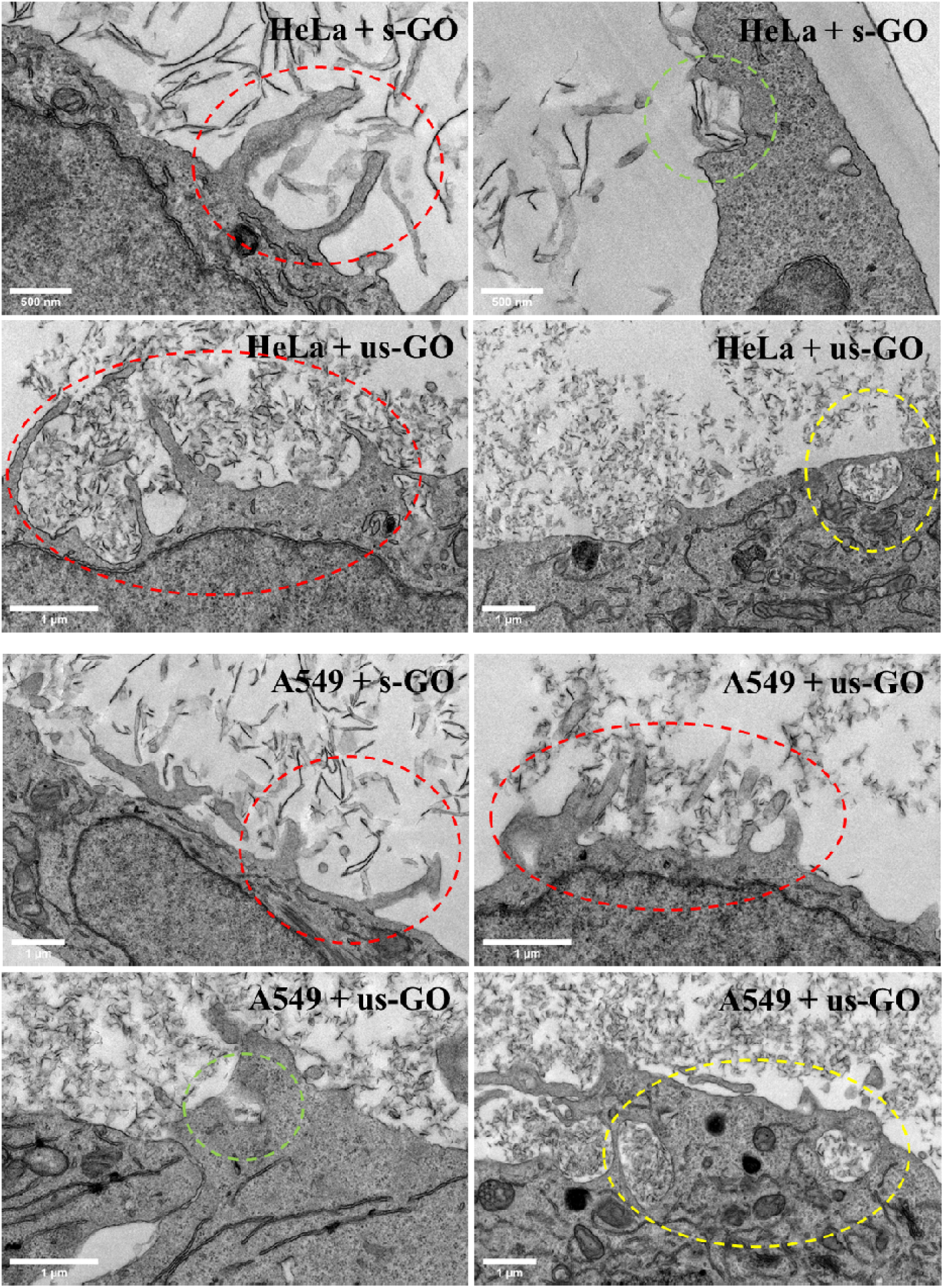
Sign of membrane ruffles (indicated by red circles) and membrane invagination (indicated by green circles) was observed in HeLa and A549 cells by TEM imaging. Also, the uptake of us-GO is found in a few cancer cells (indicated by yellow circles).

**Table S3:** Statistical analysis for the interaction of GO (s- and us-GO) with BEAS-2B, NIH/3T3, HeLa and A549 by flow cytometry (cells collected with PBS (−/−) washing); see **Figure 2e** for the corresponding graph. Tables showed the statistical analysis for the interaction of GO (s- and us-GO): **(a)** compared across the four cell lines, and **(b)** within each cell line. The data were statistically analysed using analysis of variance (two-way ANOVA) with Tukey’s multiple comparisons test. n = 3 with duplicates *Statistically different: NS = non-significant, \*\*\*\**p* < 0.0001.

| (a) | s-GO | us-GO |
| --- | --- | --- |
| HeLa vs A549 | **** | **** |
| HeLa vs NIH/3T3 |  |  |
| HeLa vs BEAS-2B |  |  |
| A549 vs NIH/3T3 | NS | NS |
| A549 vs BEAS-2B |  |  |
| NIH/3T3 vs BEAS-2B |  |  |

| (b) | HeLa | A549 | NIH/3T3 | BEAS-2B |
| --- | --- | --- | --- | --- |
| s-GO vs us-GO | **** |  |  |  |

| (b) | HeLa | A549 | NIH/3T3 | BEAS-2B |
| --- | --- | --- | --- | --- |
| s-GO vs us-GO | **** |  |  |  |

**Figure S17:**
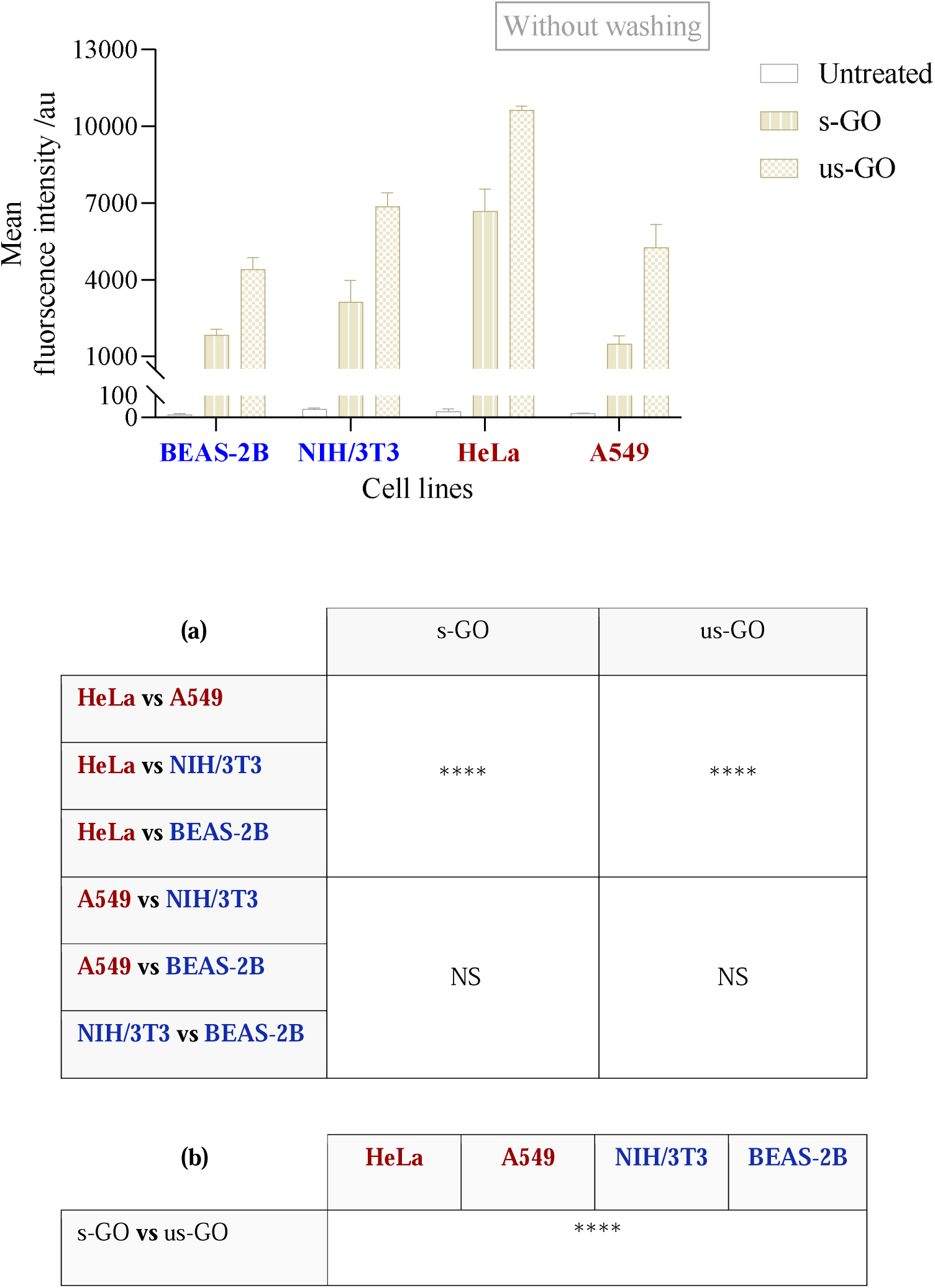
Interaction of GO (s- and us-GO) with HeLa, A549, BEAS-2B and NIH/3T3 cells by flow cytometry (cells were collected without PBS (−/−) washing). Tables showed the statistical analysis for the interaction of GO **(a)** compared across the four cell lines and **(b)** within each cell line. The data were statistically analysed using analysis of variance (two-way ANOVA) with Tukey’s multiple comparisons test. n = 3 with duplicates *Statistically different: NS = non-significant, \*\*\*\**p* < 0.0001.

**Figure S18:**
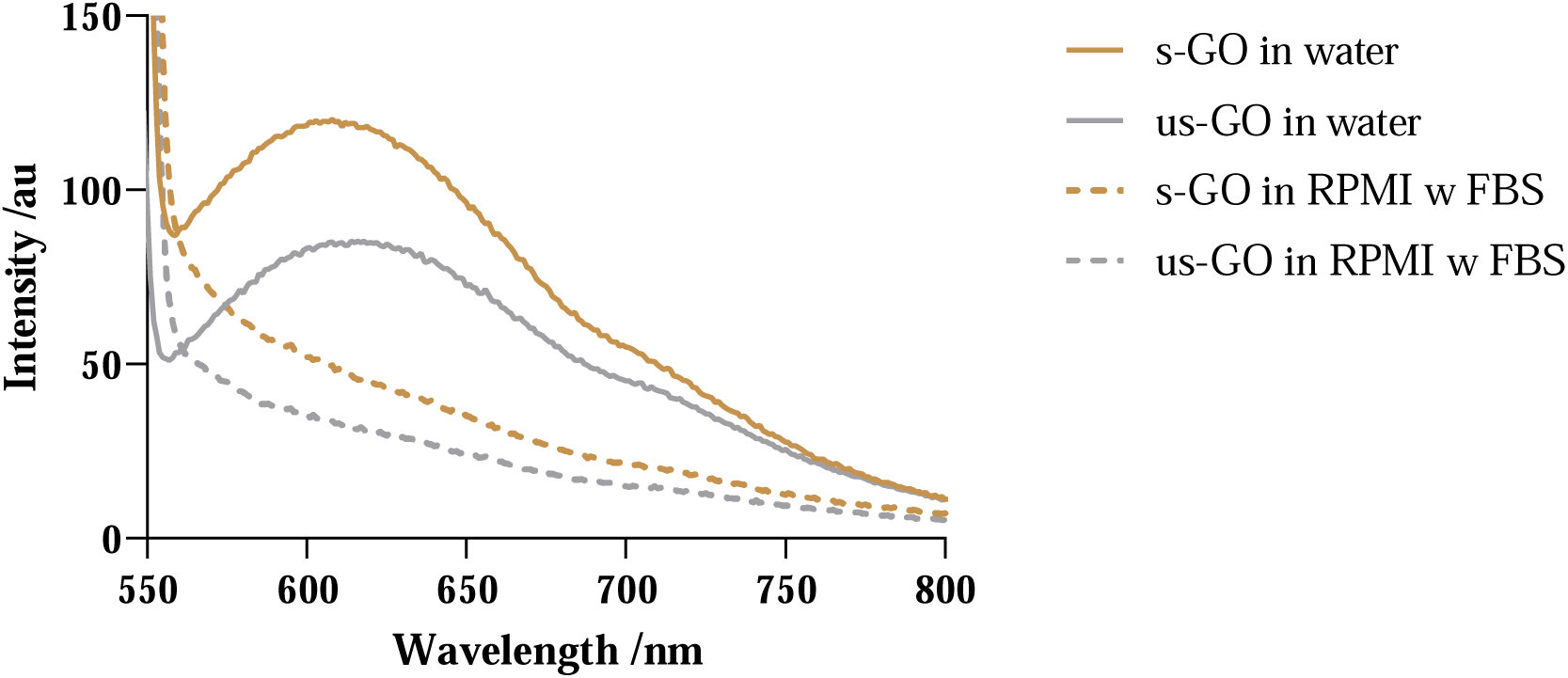
Emission spectra of s-GO and us-GO (2 mg/mL) in water (solid lines) and RPMI with 10 % FBS (materials freshly prepared in complete medium and re-suspended in water for measurement). Excitation wavelength of 525 nm is used.

**Figure S19:**
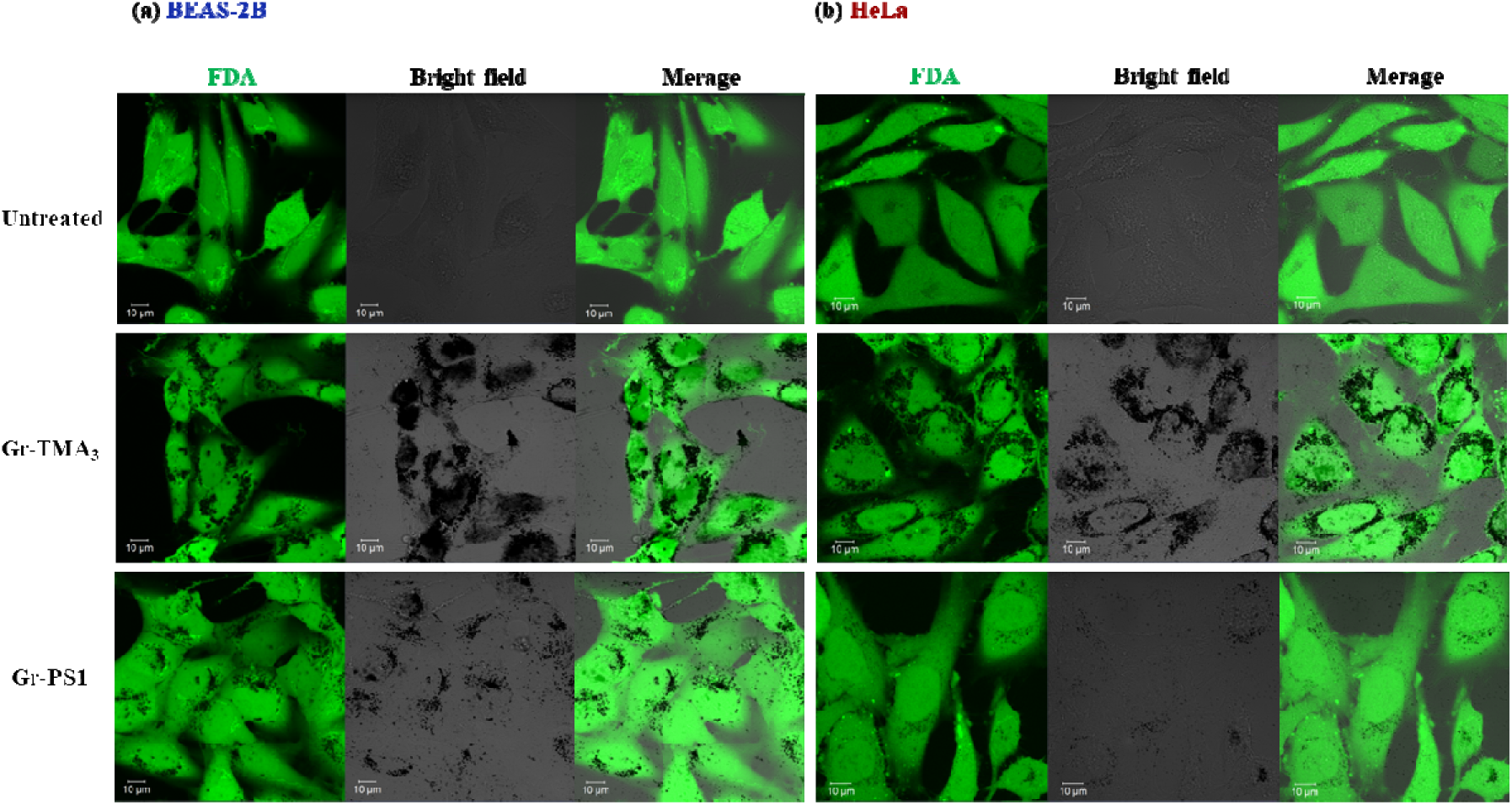
Interaction of Gr-TMA_3_ and Gr-PS1 (50 μg/mL, 24 h) in **(a)** BEAS-2B and **(b)** HeLa cells by confocal imaging. Green = FDA dye labelled cells, black = graphene flake.

**Figure S20:**
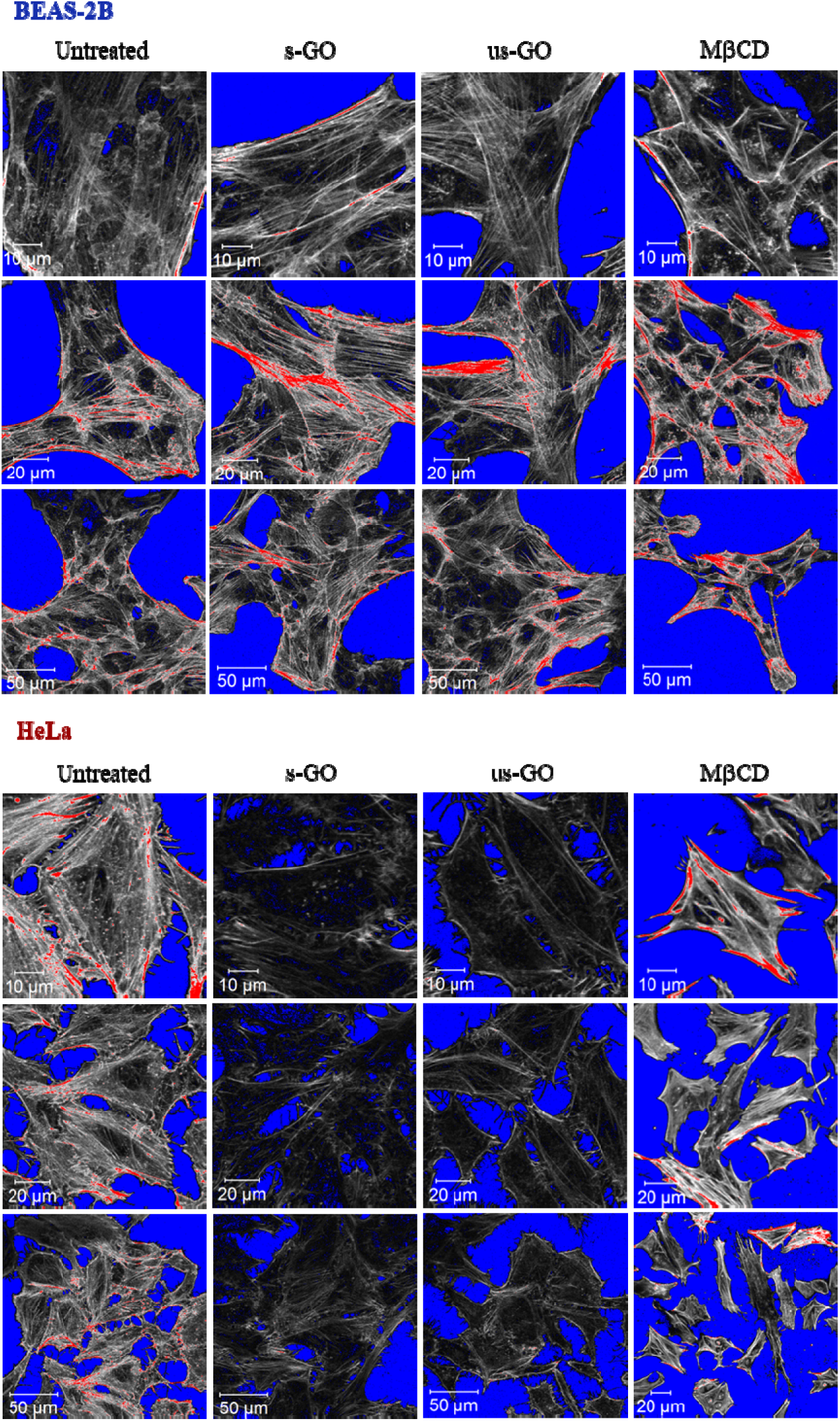
Actin filament staining for BEAS-2B and HeLa cells with and without treatment of GO (s-GO or us-GO, 100 μg/mL, 24 h) or M βCD (5 mM, 4 h). Images are shown in range indicator (pixels with zero, low, high, and maximum intensity are shown in blue, black, white and red, respectively).

**Figure S21:**
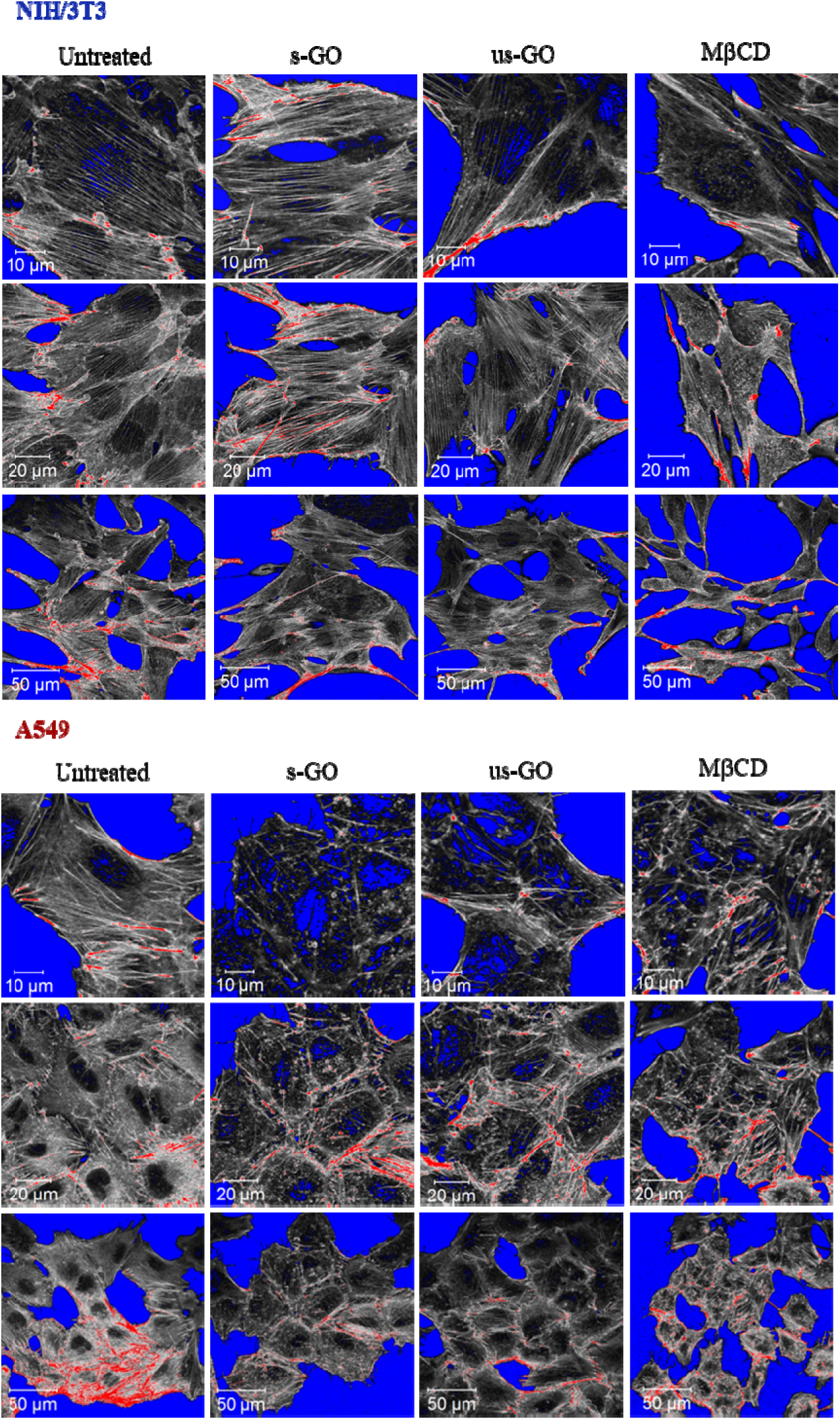
Actin filament staining for NIH/3T3 and A549 cells with and without treatment of GO (s-GO or us-GO, 100 μg/mL, 24 h) or M βCD (5 mM, 4 h). Images are shown in range indicator (pixels with zero, low, high, and maximum intensity are shown in blue, black, white and red, respectively).

